# AMPK reinforces mitochondrial metabolism and suppresses pathological remodeling in Complex V–deficient cardiomyocytes

**DOI:** 10.64898/2026.09.14.751384

**Authors:** Esteban Palacios-Contreras, Karen an der Brügge, Jakob Fell, Till Stephan, Hugo Amedei, Christof Lenz, Jelena Pesek, R. Verena Taudte, Felix Lange, Stefan Jakobs, Samuel Sossalla, Laura C. Zelarayán, Alfredo Cabrera-Orefice, Lukas Cyganek, Mario G. Pavez-Giani

## Abstract

*TMEM70* variants represent the most common nuclear cause of mitochondrial ATP synthase (Complex V) deficiency and are associated with particularly severe cardiac manifestations. Yet, how *TMEM70* deficiency disrupts cardiomyocyte metabolic maturation and function remains poorly understood, in part because suitable human disease models are lacking. Here, we model *TMEM70*-related Complex V deficiency using CRISPR-engineered human induced pluripotent stem cells differentiated into cardiomyocytes. While *TMEM70*-deficient pluripotent cells retain mitochondrial function, differentiated cardiomyocytes develop reduced mitochondrial membrane potential, impaired respiratory capacity, and pathological remodeling, revealing a differentiation-dependent failure of metabolic maturation. Chronic activation of AMP-activated protein kinase (AMPK) restores mitochondrial respiratory capacity despite persistent Complex V deficiency. Proteomic and metabolomic analyses reveal that AMPK activation in *TMEM70*-deficient cardiomyocytes reinforces mitochondrial and fatty-acid metabolism while suppressing pathological structural remodeling, accompanied by improved cardiac function. Together, these findings identify AMPK-dependent metabolic remodeling as a state-dependent mechanism of functional rescue in mitochondrial cardiomyopathy.

## Main

Complex V or ATP synthase deficiency is a primary mitochondrial disorder that impairs ATP production, disrupting cellular energy homeostasis^1,2^. Because cardiac muscle contraction relies heavily on a constant and efficient energy supply, the heart is the organ most vulnerable to mitochondrial dysfunction^3,4^. Cardiac manifestations such as hypertrophic cardiomyopathy, arrhythmogenic episodes, and heart failure are common in patients with Complex V deficiency^5^. TMEM70 is an essential assembly factor for mitochondrial Complex V^6–8^, while pathogenic variants in *TMEM70* cause Complex V deficiency, a devastating condition characterized by profound energy deficiency, multi-organ involvement, and a severe cardiac phenotype^9,10^. These patients frequently exhibit significantly diminished life expectancy early in life, and currently, there are no effective disease-modifying treatments available^11,12^. Despite its clinical severity, the molecular and physiological mechanisms by which *TMEM70* variants disrupt cardiac function and organ development remain poorly understood. This knowledge gap largely reflects the lack of suitable experimental animal models, as *Tmem70* knockout is embryonically lethal in mice^13^, impeding faithful recapitulation of the cardiometabolic and electrophysiological abnormalities observed in patients. Developing physiologically relevant disease models is therefore essential to elucidate the underlying mechanisms and enable the identification of targeted therapies for *TMEM70*-associated cardiomyopathy and related conditions.

To better understand the effects of *TMEM70* deficiency on cardiac physiology, we have created mono- and biallelic *TMEM70* loss-of-function iPSC lines through CRISPR-Cas9 genome editing, using a well-established wildtype (WT) iPSC line (Fig. 1A-B; Extended data Fig. 1A-C). Frameshift variants were introduced by targeting *TMEM70* exon 1, and the clinically predominant splice-site variant c.317-2A>G was precisely inserted into intron 2 to recapitulate the previously described patient genotype and phenotype^10^. The established iPSC lines, TMEM70^KO/WT^, TMEM70^KO/KO^, TMEM70^SNP/WT^ and TMEM70^SNP/SNP^, were verified for genome stability and pluripotent markers after editing (Extended data Fig. 1D-G). Given that the biallelic loss of *TMEM70* results in a clear cardioneuromuscular phenotype in patients, we first assessed baseline mitochondrial function in these undifferentiated iPSCs. The TMEM70^KO/KO^ iPSCs demonstrated a normal mitochondrial membrane potential similar to TMEM70^WT/WT^ cells; whereas TMEM70^SNP/SNP^ cells showed a slight increase in mitochondrial membrane potential (Extended data Fig. 2A). Both *TMEM70*-deficient lines displayed a modest increase in maximal mitochondrial respiration, while basal, spare, and ATP-linked respiration remained unchanged (Extended data Fig. 2B-C). In addition, confluence analysis revealed a marked reduction in the proliferative capacity of both *TMEM70*-deficient iPSC lines (Extended data Fig. 2D-E).

**Figure 1.**
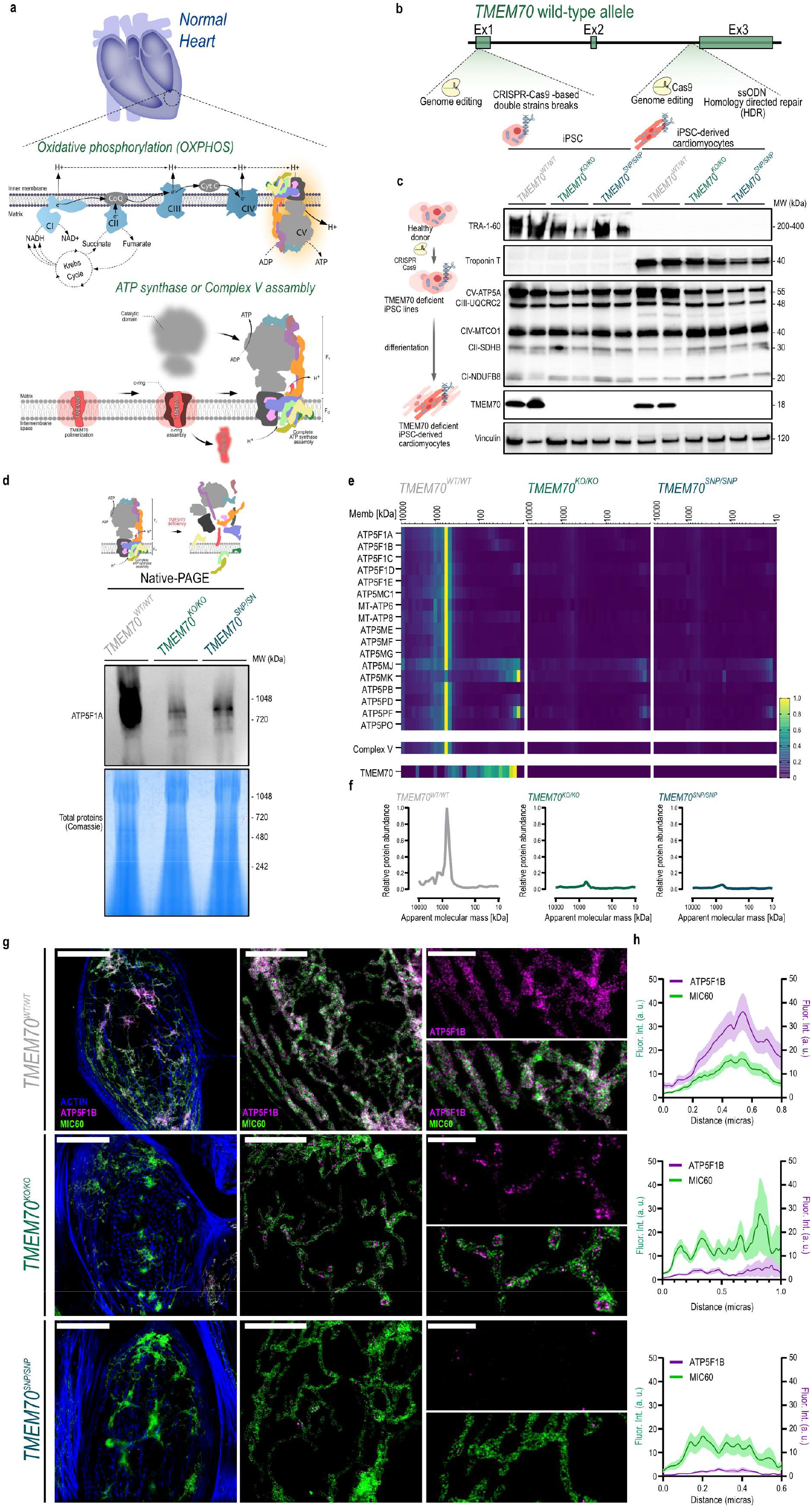
Generation and characterization of *TMEM70*-deficient iPSC-derived cardiomyocytes. (A) Schematic representation of oxidative phosphorylation and TMEM70 function in mitochondrial ATP synthase (Complex V) assembly. (B) CRISPR-Cas9 editing strategies used to generate *TMEM70*-deficient iPSC lines by targeting exon 1 (TMEM70^KO/KO^) or introducing the c.317-2A>G splice-site variant in intron 2 (TMEM70^SNP/SNP^). (C) Validation of TMEM70 expression in TMEM70^WT/WT^, TMEM70^KO/KO^) and TMEM70^SNP/SNP^ lines at the iPSC and iPSC-derived cardiomyocyte (iPSC-CM) stages. Immunoblot analysis shows TMEM70 protein levels together with stage-specific markers TRA-1-60 (for iPSCs) and cardiac troponin T (for cardiomyocytes) and representative OXPHOS subunits of Complex I–V. Vinculin served as loading control. (D) Blue native PAGE analysis of ATP synthase (Complex V) assembly in TMEM70^WT/WT^, TMEM70^KO/KO^ and TMEM70^SNP/SNP^ iPSC-CMs. (E) Complexome profiling of Complex V assembly in *TMEM70*-deficient iPSC-CMs. Heatmaps show the relative abundance and distribution of ATP synthase subunits across assembly states. (F) Complex V assembly profiles highlighting alterations in assembly intermediates in *TMEM70*-deficient iPSC-CMs compared with TMEM70^WT/WT^. (G) Dual-color STED microscopy of TMEM70^WT/WT^, TMEM70^KO/KO^ and TMEM70^SNP/SNP^ iPSC-CMs labeled for MIC60 (cristae marker), ATP5F1B (Complex V subunit) and phalloidin (F-actin). Scale bar = 20, 5 and 2 µm, respectively. (H) Quantification of mitochondrial MIC60 and ATP5F1B fluorescence intensity along mitochondrial segments. Line-scan analysis was performed across ten independent segments per condition.

Differentiation into cardiomyocytes was confirmed by a robust expression of cardiac markers, including cardiac troponin T (TNNT2), alongside loss of pluripotency markers such as TRA-1-60 (Fig. 1C). These iPSC-CMs were then analyzed to determine the impact of *TMEM70* deficiency. Biallelic loss of *TMEM70* severely impaired ATP synthase assembly (Fig. 1D) and disrupted the mitochondrial ultrastructure (Extended data Fig. 3A-B). Specifically, electron microscopy revealed disorganized cristae and altered mitochondrial morphology in *TMEM70*-deficient cells, compared to the well-organized mitochondrial network in control iPSC-CMs (Extended data Fig. 3A-B). Complexome profiling confirmed a marked defect in Complex V assembly in *TMEM70*-deficient cardiomyocytes (Fig. 1E and F), whereas the assembly of other electron transport chain (ETC) complexes remained largely preserved (Extended data Fig. 3C-E). Residual complex V holoenzyme remained detectable, consistent with previous evidence that TMEM242 acts as a partially redundant c8-ring assembly factor that cooperates with TMEM70 during ATP synthase biogenesis, potentially allowing partial compensation^14^. As documented previously^15^, super-resolution STED microscopy further revealed altered cristae organization, reduced mitochondrial ATP5F1B abundance, and decreased co-localization of ATP5F1B with MIC60 in *TMEM70*-deficient cells (Fig. 1G and H).

To assess the functional consequences of these structural defects, we next examined mitochondrial bioenergetics. *TMEM70*-deficient cardiomyocytes exhibited reduced mitochondrial membrane potential (Fig. 2A-C) and impaired respiratory capacity, with reduced basal, ATP-linked, and maximal respiration (Fig. 2D-E). Consistently, proteomic profiling revealed reduced abundance of mitochondrial metabolic proteins and increased expression of proteins associated with pathological cardiac remodeling (Fig. 2F-K). Functionally, *TMEM70*-deficient cardiomyocytes exhibited impaired contractile performance, reflected by reduced contraction velocity and amplitude, together with an increased spontaneous beating rate (Fig. 2L and M). These functional abnormalities were accompanied by broad downregulation of sarcomeric and Ca^2+^ handling genes (Fig. 2N).

**Figure 2.**
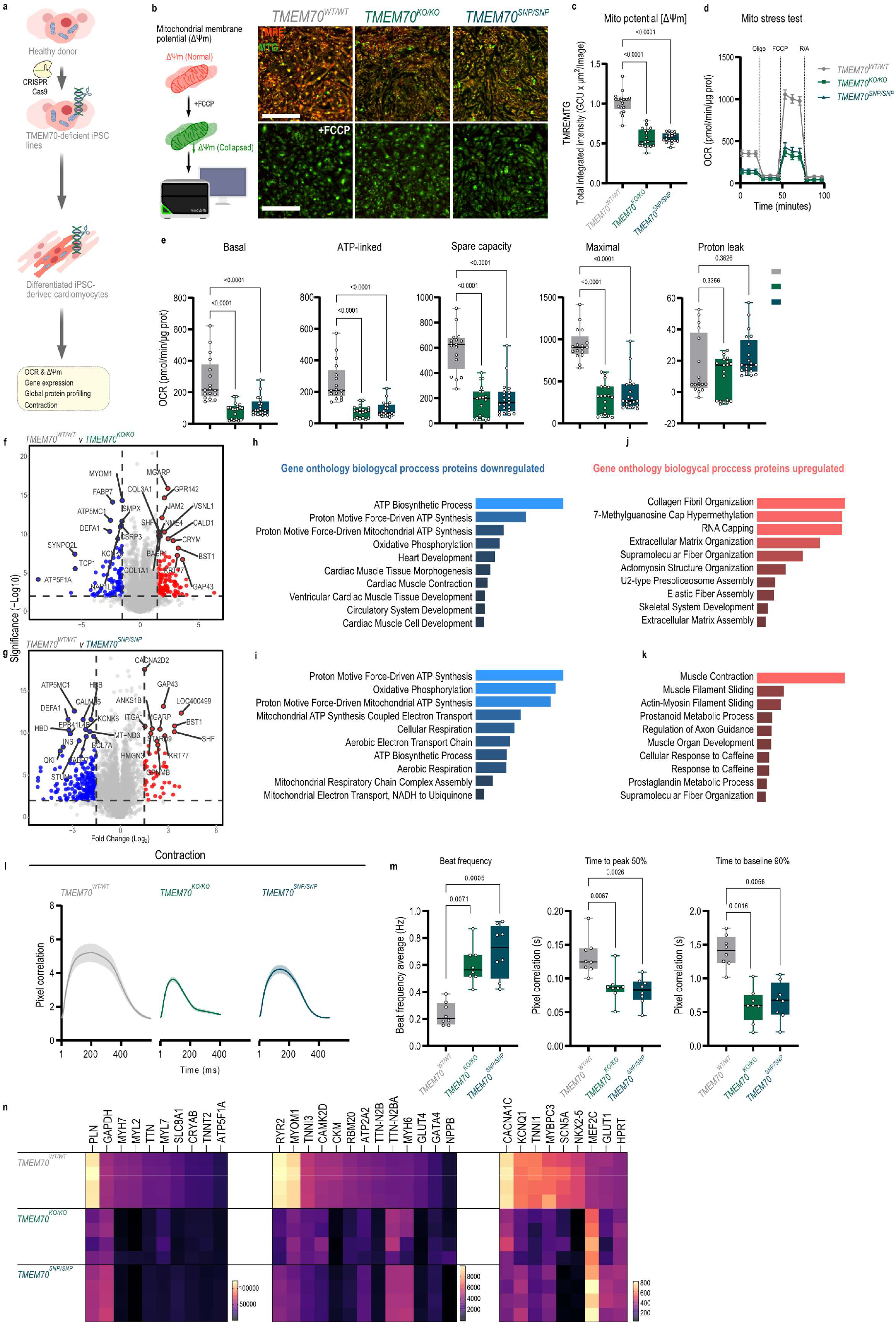
*TMEM70* deficiency impairs mitochondrial function, metabolic remodeling and contractile performance in iPSC-derived cardiomyocytes. (A) Schematic overview of cardiomyocyte differentiation and experimental workflow. TMEM70^WT/WT^, TMEM70^KO/KO^ and TMEM70^SNP/SNP^ iPSCs were differentiated into iPSC-CMs and analyzed at day 65 for mitochondrial function, proteomic remodeling, and contractile performance. (B) Representative fluorescence images of mitochondrial membrane potential assessed using TMRE, with MitoTracker Green (MTG) as a mitochondrial mass control. FCCP treatment was used as a depolarization control. Scale bar = 100 µm. (C) Mitochondrial membrane potential (Δψm) as TMRE/MTG total integrated intensity in TMEM70^WT/WT^, TMEM70^KO/KO^, and TMEM70^SNP/SNP^ iPSC-CMs; n=18 from three independent differentiations. P values were determined by one-way ANOVA with Tukey’s multiple comparisons test. (D) Oxygen consumption rate (OCR) during mitochondrial stress test for TMEM70^WT/WT^, TMEM70^KO/KO^, and TMEM70^SNP/SNP^ iPSC-CMs with sequential addition of oligomycin, FCCP and rotenone/antimycin A. (E) Quantification of basal, ATP-linked, maximal, spare respiratory capacity and proton leak, normalized to total protein; n=17-21 from three independent differentiations. P values were determined by Kruskal-Wallis test. (F–G) Volcano plots of global proteome profiles of TMEM70^KO/KO^ and TMEM70^SNP/SNP^ iPSC-CMs compared with TMEM70^WT/WT^; n=5 from two independent differentiations. (H–K) Gene ontology biological process enrichment analysis of differentially expressed proteins decreased (H,I) or increased (J,K) in *TMEM70*-deficient iPSC-CMs. (L) Representative contraction traces illustrating altered contraction kinetics in TMEM70^WT/WT^, TMEM70^KO/KO^, and TMEM70^SNP/SNP^ iPSC-CMs. (M) Quantification of spontaneous beat frequency, time to 50% peak of contraction, and time to 90% return to baseline in TMEM70^WT/WT^, TMEM70^KO/KO^, and TMEM70^SNP/SNP^ iPSC-CMs; n=8 from four independent differentiations. P values were determined by Kruskal-Wallis test. (N) NanoString gene expression analysis showing altered expression of cardiac and metabolic genes in TMEM70^WT/WT^ and *TMEM70*-deficient iPSC-CMs; n=4 from three independent differentiations.

Consistent with clinical observations in heterozygous carriers, monoallelic loss of *TMEM70* (TMEM70^KO/WT^ and TMEM70^SNP/WT^) did not reduce Complex V protein abundance, as ATP5F1A levels remained comparable to TMEM70^WT/WT^ (Extended Data Fig. 4A-B). Mitochondrial respiratory function was likewise unaffected in monoallelic *TMEM70* mutant lines, with basal, ATP-linked, maximal, and spare respiratory capacities comparable to TMEM70^WT/WT^ cells (Extended Data Fig. 4C-F). These findings indicate that a single functional *TMEM70* allele suffices to preserve Complex V abundance and overall oxidative phosphorylation capacity in cardiac cells.

Whereas biallelic *TMEM70* loss had only minor effects on mitochondrial oxygen consumption in undifferentiated iPSCs, pronounced mitochondrial dysfunction emerged following cardiomyocyte differentiation, suggesting that TMEM70 becomes increasingly critical as cardiomyocytes acquire a more oxidative metabolic state. We therefore compared mitochondrial respiration in early day-35 and more mature day-65 cardiomyocytes. At day 35, *TMEM70*-deficient cardiomyocytes exhibited respiratory capacities comparable to TMEM70^WT/WT^ cells, whereas by day 65 respiratory capacity had markedly increased in WT cardiomyocytes, while TMEM70^KO/KO^ and TMEM70^SNP/SNP^ iPSC-CM cultures remained metabolically static (Extended data Fig. 5B-D), indicating impaired acquisition of oxidative capacity during maturation. Proteome profiling further supported this maturation defect. Between days 35 and 65, TMEM70^WT/WT^ cardiomyocytes showed increased abundance of ETC-associated proteins and reduced abundance of glycolytic enzymes, including LDHA, ENO1 and PGK1, consistent with the transition from glycolytic to oxidative energy metabolism (Extended Data Fig. 5E and G). In *TMEM70*-deficient cells, maturation-associated changes in fatty-acid oxidation proteins were attenuated, whereas extracellular matrix and cytoskeletal proteins underwent more pronounced remodeling than in WT cells, revealing a divergence between metabolic maturation and structural remodeling (Extended Data Fig. 5F-H).

Using this maturation window, *TMEM70*-deficient cardiomyocytes were treated for 30 days with three compounds previously reported to enhance mitochondrial oxidative capacity and promote metabolic maturation: ZLN005, an activator of peroxisome proliferator-activated receptor-γ coactivator 1α (PGC-1α)^16^; β-hydroxybutyrate, which promotes ketone body metabolism^17^; and AMPK activator MK8722^18,19^ (Fig. 3A). Among the tested conditions, only MK8722 at 1 µM significantly increased mitochondrial membrane potential (Fig. 3B). Dose-response analysis confirmed that MK8722 did not adversely affect cardiomyocyte viability at the concentration used (Extended data Fig. 6A). Prolonged MK8722 treatment markedly enhanced mitochondrial respiratory capacity in *TMEM70*-deficient iPSC-CMs, with increased maximal respiration and spare respiratory capacity (Fig. 3C-D). Notably, basal and ATP-linked respiration were reduced, while proton leak remained unchanged, suggesting a shift toward lower basal energetic demand while preserving an enhanced respiratory reserve (Fig. 3D). These improvements in mitochondrial function were accompanied by elevated levels of OXPHOS proteins and key regulators of mitochondrial dynamics, including mitofusin 1 (MFN1), mitofusin 2 (MFN2), and optic atrophy protein 1 (OPA1), while mitochondrial outer membrane marker TOMM20 remained unchanged (Fig. 3E-F). Native complexome profiling of the mitochondrial fraction further revealed that AMPK activation increased the abundance of mature Complex II, with a mild enhancement of Complex V assembly (Fig. 3G and H), without major changes in the assembly of other ETC complexes (Extended data Fig. 6B).

**Figure 3.**
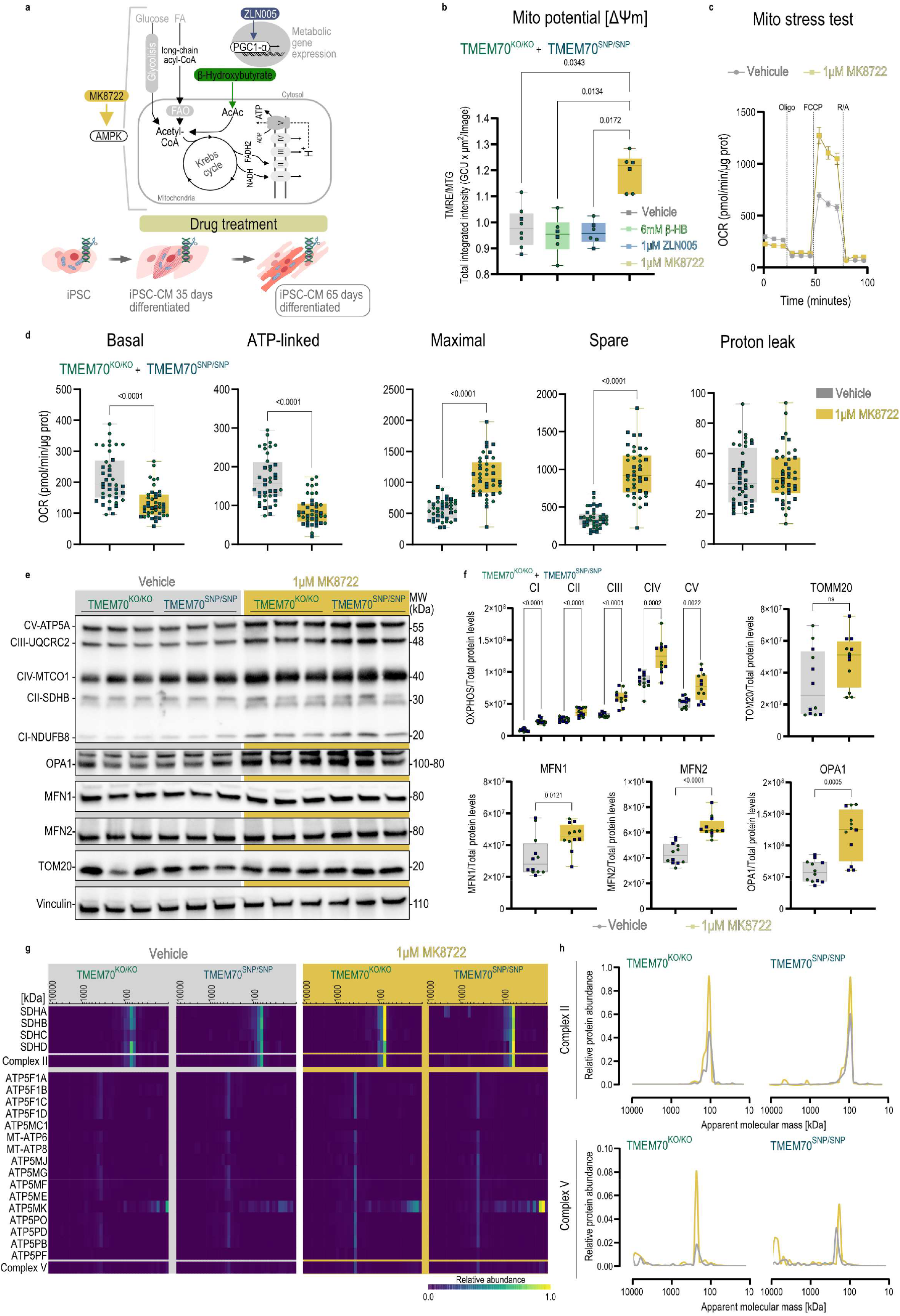
AMPK activation restores mitochondrial respiratory capacity in *TMEM70*-deficient cardiomyocytes. (A) Schematic overview of metabolic interventions during cardiomyocyte maturation. *TMEM70*-deficient iPSC-CMs were treated for 30 days (day 35 to day 65) with the PGC1α activator ZLN005 (1 µM), the ketone body β-hydroxybutyrate (6 mM), or the pan-AMPK activator MK8722 (1 µM). (B) Mitochondrial membrane potential (Δψm) as TMRE/MTG total integrated intensity in *TMEM70*-deficient iPSC-CMs (TMEM70^KO/KO^ and TMEM70^SNP/SNP^) following treatment with vehicle (DMSO), β-hydroxybutyrate, ZLN005, or MK8722; n=6-8 from three independent differentiations. P values were determined by Kruskal-Wallis test. (C) Oxygen consumption rate during mitochondrial stress test in *TMEM70*-deficient iPSC-CMs (TMEM70^KO/KO^ and TMEM70^SNP/SNP^), following vehicle (DMSO) and MK8722 treatment; n=40 from four independent differentiations. (D) Quantification of basal, ATP-linked, maximal, spare respiratory capacity, and proton leak, normalized to total protein; n=40 from four independent differentiations. P values were determined by two-tailed Mann Whitney test. (E) Representative immunoblot analysis of OXPHOS proteins, OPA1, MFN1, MFN2, and TOMM20 in vehicle- and MK8722-treated TMEM70^KO/KO^ and TMEM70^SNP/SNP^ iPSC-CMs. Vinculin served as loading control. (F) Densitometric quantification of OXPHOS proteins, TOMM20, MFN1, MFN2 and OPA1 in vehicle- and MK8722-treated TMEM70^KO/KO^ and TMEM70^SNP/SNP^ iPSC-CMs, normalized to total protein; n=12 from four independent differentiations. P values were determined by two-tailed Mann Whitney test. (G) Complexome profiling of Complex II and Complex V assembly in vehicle- and MK8722-treated TMEM70^KO/KO^ and TMEM70^SNP/SNP^ iPSC-CMs. Heatmaps show the relative abundance and distribution of succinate dehydrogenase (Complex II) and ATP synthase (Complex V) subunits across assembly states. (H) Assembly profiles of Complex II and Complex V highlighting alterations in assembly intermediates following MK8722 treatment in TMEM70^KO/KO^ and TMEM70^SNP/SNP^ iPSC-CMs.

To define the metabolic basis of enhanced mitochondrial capacity following AMPK activation, we first examined canonical downstream targets of AMPK (Fig. 4A). Chronic MK8722 treatment increased phosphorylation of acetyl-CoA carboxylase (ACC Ser-79) and pyruvate dehydrogenase (PDH Ser-293) in *TMEM70*-deficient cardiomyocytes, consistent with inhibition of ACC and reduced PDH activity (Fig. 4B-C). These changes are expected to favor fatty-acid oxidation while limiting PDH-dependent entry of glucose-derived acetyl-CoA into the tricarboxylic acid (TCA) cycle. We performed U-^13^C_16_-palmitic acid and U-^13^C-glucose tracing to assess substrate utilization. MK8722 treatment increased fatty acid-derived labeling of TCA intermediates in both WT and *TMEM70*-deficient cardiomyocytes (Extended data Fig. 7A and Fig. 4D). Notably, chronic AMPK activation also increased glucose-derived labeling of TCA intermediates in *TMEM70*-deficient cardiomyocytes, which, in contrast to WT cells, exhibited elevated baseline pyruvate and lactate levels that were not further increased by MK8722 (Extended data Fig. 7B). Given the inhibitory phosphorylation of PDH, the increased glucose-supported TCA labeling is unlikely to reflect enhanced pyruvate oxidation via PDH and instead suggests engagement of alternative glucose-dependent anaplerotic pathways. In line with this metabolic rewiring, global proteomic analysis revealed upregulation of mitochondrial and fatty-acid metabolism–associated proteins upon AMPK activation, including CPT1B, CPT2, SIRT3, CD36 and ACADVL (Fig. 4E-F), together with reduced abundance of proteins linked to extracellular matrix organization and pathological cardiac remodeling, such as COL1A1, MYH7, ANKRD1, CCN2 and NFATC3 (Fig. 4G). Functionally, MK8722-treated *TMEM70*-deficient cardiomyocytes displayed normalization of spontaneous beating frequency and improved contraction kinetics (Fig. 4H). At the molecular level, AMPK activation reduced phosphorylation of both RyR2 Ser-2814, and PLN Thr-17, while CaMKII Thr-286 phosphorylation and Sarco/Endoplasmic Reticulum Calcium ATPase 2 (SERCA2) protein levels remained unchanged (Fig. 4I-J), indicating coordinated remodeling of sarcoplasmic reticulum Ca^2+^ handling.

**Figure 4.**
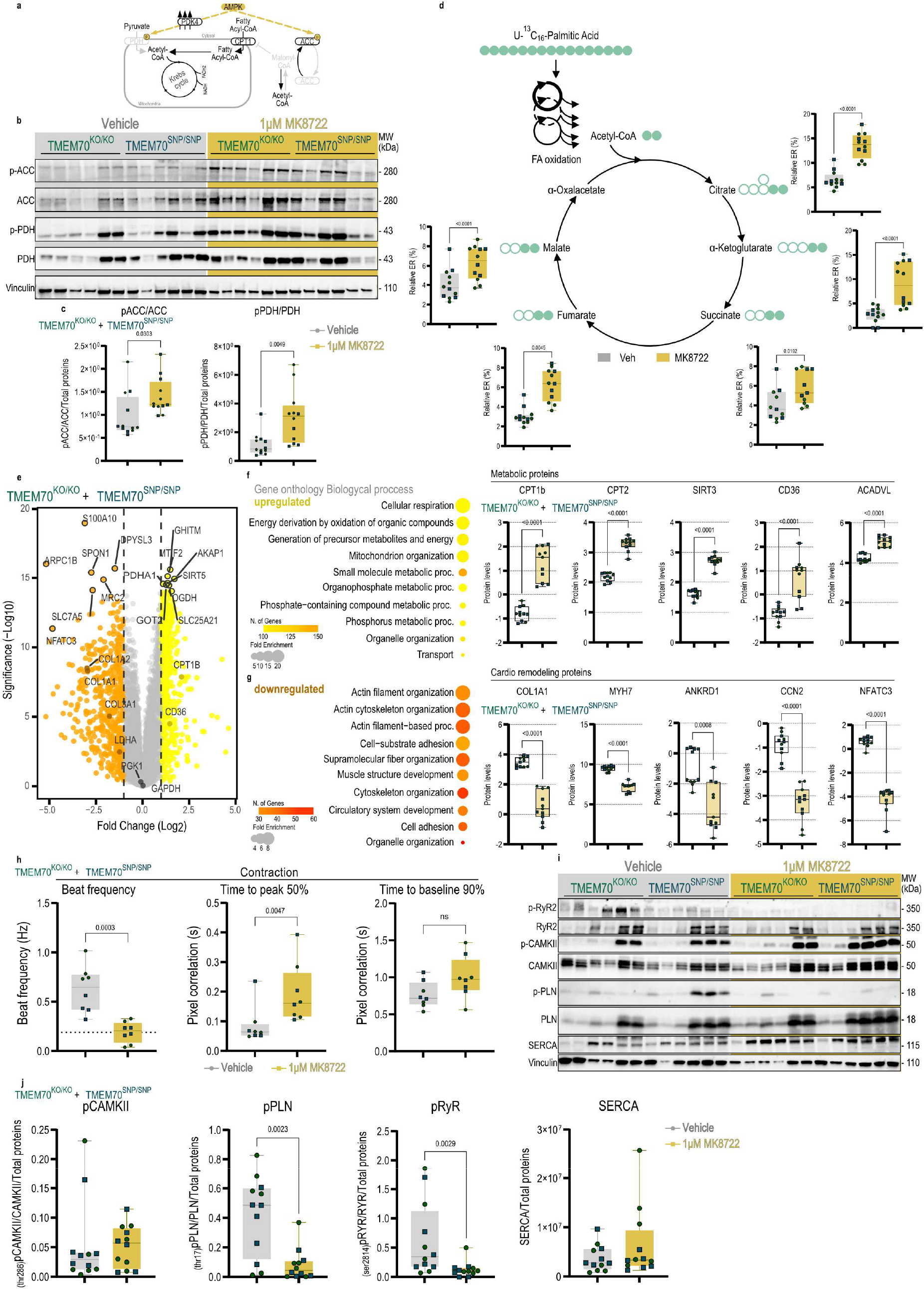
AMPK induces coordinated metabolic, structural and Ca^2+^-handling remodeling in *TMEM70*-deficient cardiomyocytes. (A) Schematic overview of AMPK-dependent regulation of fatty-acid and glucose metabolism through ACC and PDH signaling. (B) Representative immunoblots of ACC and PDH signaling activity in *TMEM70*-deficient iPSC-CMs (TMEM70^KO/KO^ and TMEM70^SNP/SNP^; blue square), after 30 days of vehicle or 1 µM MK8722 treatment. Vinculin served as loading control. (C) Quantification of phosphorylated protein levels (pACC/ACC and pPDH/PDH) in vehicle- and MK8722-treated TMEM70^KO/KO^ and TMEM70^SNP/SNP^ iPSC-CMs; n=12 from four independent differentiations. P values were determined by two-tailed Mann Whitney test. (D) U-^13^C_16_-palmitic acid tracing in vehicle- and MK8722-treated TMEM70^KO/KO^ and TMEM70^SNP/SNP^ iPSC-CMs. The schematic illustrates fatty-acid β-oxidation and entry of labeled acetyl-CoA into the TCA cycle. Relative isotopic enrichment (ER, %) of citrate, α-ketoglutarate, succinate, fumarate, and malate is shown; filled circles indicate ^13^C-labeled carbons; n=11 from four independent differentiations. P values were determined by two-tailed Mann Whitney test. (E) Volcano plots of global proteomic profiles of vehicle-versus MK8722-treated TMEM70^KO/KO^ and TMEM70^SNP/SNP^ iPSC-CMs. Data represent relative protein abundance changes across conditions; n=12 from four independent differentiations. (F-G) Gene ontology biological process enrichment analysis of proteins increased and decreased following MK8722 treatment in TMEM70^KO/KO^ and TMEM70^SNP/SNP^ iPSC-CMs. Quantification of selected proteins associated with these pathways is shown; n=12 from four independent differentiations. P values were determined by two-tailed Mann Whitney test. (H) Quantification of spontaneous beat frequency, time to 50% peak contraction and time to 90% return to baseline in vehicle- and MK8722-treated TMEM70^KO/KO^ and TMEM70^SNP/SNP^ iPSC-CMs; n=8; from four independent differentiations. P values were determined by two-tailed Mann Whitney test. (I) Representative immunoblots of Ca^2+^-handling regulators, including phosphorylated and total RyR2 (Ser2814), CaMKII, phospholamban (PLN) and SERCA2 in vehicle- and MK8722-treated TMEM70^KO/KO^ and TMEM70^SNP/SNP^ iPSC-CMs. Vinculin served as loading control. (J) Quantification of phosphorylated protein levels (pRYR2/RYR2, pPLN/PLN and pCaMKII/CaMKII) in vehicle- and MK8722-treated TMEM70^KO/KO^ and TMEM70^SNP/SNP^ iPSC-CMs; n=12 from four independent differentiations. P values were determined by two-tailed Mann Whitney test.

Defects in ATP synthase assembly profoundly alter mitochondrial bioenergetics, yet how these defects translate into cardiomyocyte dysfunction has remained unclear. Here, using a human iPSC-based model of TMEM70-related Complex V deficiency, we show that mitochondrial impairment emerges in a differentiation-dependent manner and is accompanied by a failure of coordinated metabolic and functional maturation. Chronic AMPK activation does not correct the underlying assembly defect in ATP synthase, but instead induces a coordinated remodeling program that restores mitochondrial respiratory capacity while suppressing pathological cellular remodeling. These findings define a state-dependent mechanism by which mitochondrial dysfunction can be functionally compensated through AMPK-dependent metabolic remodeling, without direct restoration of ATP synthase integrity.

A central observation of this study is that the consequences of TMEM70 deficiency are strongly dependent on cellular state. While undifferentiated iPSCs tolerate impaired ATP synthase assembly with largely preserved mitochondrial membrane potential and respiratory function, differentiated cardiomyocytes exhibit reduced Δψ_m_, impaired maximal respiration, and features of pathological remodeling. This transition coincides with the developmental shift toward oxidative metabolism and the increasing bioenergetic demands of cardiomyocyte maturation and excitation–contraction coupling^18^, suggesting that ATP synthase defects become limiting specifically as cardiomyocytes transition toward an oxidative, functionally mature state. These findings indicate that *TMEM70* deficiency primarily disrupts coordinated metabolic and structural maturation of cardiomyocytes as the underlying ATP synthase defect progresses to bioenergetic failure during maturation.

Chronic AMPK activation with MK8722 restored mitochondrial respiratory reserve in *TMEM70*-deficient cardiomyocytes and was accompanied by broad upregulation of mitochondrial proteins involved in ETC, TCA cycle and fatty-acid oxidation. This coordinated remodeling of the mitochondrial proteome was further supported by expansion of TCA intermediate pools, consistent with reinforcement of oxidative metabolic capacity rather than a simple redistribution of carbon flux. Increased phosphorylation of ACC and PDH further supports substrate remodeling toward fatty-acid metabolism, in which glucose oxidation is restrained while alternative glucose-dependent anaplerotic pathways contribute to TCA cycle metabolism^19,20^. Together, these findings support a model in which AMPK enhances mitochondrial metabolic competence through integrated regulation of substrate utilization, mitochondrial organization, and respiratory machinery in the setting of Complex V deficiency.

In parallel with mitochondrial reinforcement, AMPK activation markedly suppressed proteins associated with extracellular matrix organization, focal adhesion, and stress-associated contractile remodeling. This included multiple collagen isoforms and adhesion-associated proteins, indicating attenuation of a pro-fibrotic and mechanically stressed cellular state related to *TMEM70* deficiency. Reduced abundance of contractile and cytoskeletal proteins, including MYH7, which are implicated in hypertrophic cardiomyopathy ^21^, further supports reversal of pathological cardiac remodeling^5,22^. The preservation of core structural proteins further argues against a generalized loss of cardiomyocyte identity or maturation and instead supports a selective remodeling response to AMPK activation. Notably, decreased NFATC3 abundance provides a potential link to reduced stress-responsive transcriptional signaling^23,24^, although the underlying regulatory mechanisms remain to be defined. Together, these observations suggest that AMPK activation does not simply enhance mitochondrial metabolism, but also reshapes the broader cellular state associated with *TMEM70* deficiency. This coordinated response may explain how substantial functional recovery can occur despite persistence of the underlying ATP synthase assembly defect.

Importantly, our findings identify Ca^2+^ handling as a third mechanistic layer within the broader AMPK-induced remodeling response contributing to functional rescue. Reduced phosphorylation of RyR2 at Ser-2814 is particularly notable, as this site is a well-established CaMKII-sensitive determinant of pathological sarcoplasmic reticulum Ca^2+^ leak^25^. Together with reduced PLN Thr17 phosphorylation and unchanged bulk CaMKII phosphorylation, these findings suggest selective remodeling of sarcoplasmic reticulum Ca^2+^ handling rather than broad suppression of CaMKII signaling. This provides a potential molecular link to the normalization of beat frequency and contraction kinetics observed after treatment. More broadly, these physiological changes are consistent with coordinated normalization of cardiomyocyte state, integrating mitochondrial reinforcement, structural remodeling, and excitation–contraction coupling.

Our results highlight that Complex V–dependent mitochondrial cardiomyopathy is shaped not only by the primary respiratory defect, but also by the capacity of cardiomyocytes to adapt during maturation. The limiting factor during cardiomyocyte maturation may therefore not be ATP deficiency alone, but the ability of mitochondria to maintain sufficient electrochemical and respiratory capacity. In this context, AMPK emerges as a key regulator of this adaptive response and a potential therapeutic target for cardiac manifestations of primary mitochondrial disease.

## Methods

### Generation of *TMEM70*-deficient iPSC generation

Human iPSC lines from a healthy donor and a CRISPR/Cas9-engineered *TMEM70*-deficient line were used in this study. The study was approved by the Ethics Committee of the Justus Liebig University Giessen (approval number: 258/16), and University Medical Center Göttingen (approval number: 10/9/15) and performed in accordance with the approved guidelines. WT iPSC line UMGi014-C clone 14 (isWT1.14) was generated from dermal fibroblasts using the integration-free Sendai virus. *TMEM70* loss-of-function iPSC lines were generated using ribonucleoprotein-based CRISPR–Cas9 genome editing, as described previously^26^. For the TMEM70 knockout, exon 1 editing generated the heterozygous clone UMGi014-C-28 clone 19 (isWT1-TMEM70-Ex1-KO.19; TMEM70^KO/WT^) and the compound-heterozygous clone UMGi014-C-28 clone 17 (isWT1-TMEM70-Ex1-KO.17; TMEM70^KO/KO^). For the c.317-2A>G splice-site variant, homology-directed repair generated the heterozygous clone UMGi014-C-29 clone 6 (isWT1-TMEM70-In2-KO.6; TMEM70^SNP/WT^) and the homozygous clone UMGi014-C-29 clone 3 (isWT1-TMEM70-In2-KO.3; TMEM70^SNP/SNP^). CRISPR-modified iPSCs underwent pluripotency characterization by immunostaining and flow cytometry analysis as well as chromosomal integrity by digital karyotyping. Detailed genome editing and culture procedures are provided in the Extended Methods.

### iPSC-derived cardiomyocyte differentiation

Cardiomyocyte differentiation was performed using biphasic temporal modulation of Wnt signaling, as described previously^27^. Briefly, confluent hiPSC cultures were treated with 4 µM CHIR99021 (GSK3β inhibitor) for mesoderm induction, followed by 5 µM IWP2 (Wnt inhibition) to promote cardiac specification. Beating cardiomyocytes typically emerged between days 9–11 of differentiation. Cells were metabolically purified using glucose-depleted, lactate-containing medium for seven days. Experiments were conducted between differentiation days 35–65 unless otherwise indicated. For chronic drug treatment, chemical compounds were added to culture medium for 30 days, with media replacement every 48 h; vehicle-treated cultures [DMSO] served as controls.

### Native-PAGE, SDS-PAGE and immunoblot

Whole cells and mitochondria isolated by differential centrifugation were solubilized under native conditions and separated by Blue Native PAGE. Whole cells were solubilized in NP40-containing buffer, while mitochondria were solubilized in digitonin-containing buffer and separated on 3–12% acrylamide gradient gels. Complex V assembly was analyzed by immunoblotting using antibodies against ATP synthase subunits; ATP5F1A. For SDS-PAGE, total protein lysates were prepared in 1% Sodium deoxycholate [SDC] supplemented with protease and phosphatase inhibitors. Equal amounts of protein were separated by SDS-PAGE and transferred to PVDF membranes. Membranes were incubated with primary antibodies overnight at 4 °C, followed by HRP-conjugated secondary antibodies. Bands were visualized by chemiluminescence and quantified using Image Lab (Bio-Rad) software.

### Complexome profiling

Whole cells or isolated mitochondria were solubilized under 1% NP40 or digitonin (6 mg/mg protein), respectively, to preserve protein complexes. Protein samples were separated by Blue Native PAGE according to Wittig et al. 2006^28^, and gel lanes were sliced into 35 or 40 fractions, respectively, followed by in-gel trypsin digestion and analysis by liquid chromatography– tandem mass spectrometry (LC–MS/MS). MS data were processed using MaxQuant for protein identification and quantification. The list of identified proteins was hierarchically clustered based on Pearson correlation. Migration profiles of the identified protein groups were reconstructed to show abundance (absolute intensity) against apparent molecular mass, focusing on the assembly states of the oxidative phosphorylation (OXPHOS) complexes and their subunits. Relative abundance of mature and assembly-intermediate forms was calculated based on the maximal intensity of each protein group across conditions.

### Mitochondrial respiration assay

Oxygen consumption rate (OCR) was measured using a Seahorse XFe24 Analyzer. iPSC-derived cardiomyocytes were seeded onto Matrigel-coated XFe24 plates seven days prior to the assay. Cartridge sensors were equilibrated in calibrant assay medium 24 h prior to measurement. Basal respiration was recorded, followed by sequential injection of oligomycin (ATP synthase inhibitor; [1µM]), FCCP (uncoupler; [1µM]), and rotenone/antimycin A (complex I/III inhibitors; [0.5 and 0.5µM, respectively]). ATP-linked respiration was calculated as the difference between basal OCR and oligomycin-treated OCR. Maximal respiration was defined as FCCP-stimulated OCR. Spare respiratory capacity was calculated as maximal minus basal respiration. Proton leak was defined as OCR remaining after oligomycin treatment minus non-mitochondrial respiration. OCR values were normalized to protein concentration using BCA methods.

### Mitochondrial potential assay

Mitochondrial membrane potential (Δψm) was assessed using tetramethylrhodamine methyl ester (TMRM) and MitoTracker Green (MTG) for mitochondrial mass normalization. Cells were cultured with dye in non-quenching mode [20 nM] and visualized using an Incucyte automated-imaging system. Mean TMRM (red channel) fluorescence intensity per cell was quantified and normalized to MTG (green channel) signal. For depolarization controls, 5 µM FCCP was applied acutely.

### STED microscopy

Super-resolution stimulated emission depletion (STED) microscopy was performed to assess mitochondrial network organization and cristae-associated structures. Cells were fixed in 8% paraformaldehyde, permeabilized, and stained with Phalloidin-OregonGreen (1:100), M292 (1:100, MIC60, rabbit) and M346 (1:200, ATP5F1B, mouse) for 1.5 h at room temperature. Secondary antibodies conjugated to STED-compatible fluorophores anti-rb-StarRed and anti-ms-Alexa594 were applied for 1 h at room temperature. Images were acquired using a STED microscope Abberior Expert Line dual-color STED. Data were smoothed with a low-pass filter in Imspector. Mitochondrial morphology metrics were quantified using ImageJ.

### NanoString

Total RNA was extracted using column-based purification. RNA quality and concentration were assessed prior to hybridization. Gene expression profiling was performed using NanoString nCounter technology with a customized panel targeting mitochondrial and cardiac genes. Raw counts were normalized to internal housekeeping controls and analyzed using nSolver software.

### Proteome profiling

Whole-cell lysates were denatured, reduced, and alkylated followed by trypsin digestion using the iST kit (PreOmics). The resulting peptides were separated and analyzed by LC–MS/MS in data-independent acquisition (DIA) mode. Protein identification and quantification were performed using label-free quantification in DIA-NN v2.2. Differentially expressed proteins were identified using statistical thresholds adjusted for pairwise or multiple testing.

### Contractility measurements in spontaneously beating hiPSC-CMs

Spontaneously beating cardiomyocytes were recorded using phase-contrast video microscopy (IonOptix). Contractile parameters, including beating frequency, contraction amplitude, and time to 50% peak contraction, were analyzed using motion-tracking software. At least two independent differentiations were analyzed per condition.

### Cytotoxicity assay

Cell viability following compound treatment was assessed using fluorescence-based Calcein-AM and propidium iodide staining. Cells were exposed to increasing concentrations of MK8722, and viability was normalized to untreated controls. Measurements were performed in triplicate.

### Carbon tracing and metabolomic analysis

For carbon tracing experiments, hiPSC-derived cardiomyocytes were incubated in either glucose-free RPMI supplemented with B27, 2 mM L-glutamine, and [U-^13^C_6_]glucose [10 mM] or RPMI supplemented with B27, 2 mM L-glutamine, carnitine [0.5mM], and [U-^13^C_6_]Palmitic-acid [100 µM] conjugated with BSA [1:3 ratio]. MK8722 [1 µM] was maintained throughout the labeling period. Cells were labeled for 48 h under standard culture conditions. Metabolism was quenched by rapid aspiration of medium followed by immediate addition of ice-cold 80% methanol. Cells were incubated on dry ice, scraped into extraction solvent, and centrifuged at 16,000g for 10 min at 4 °C. Extracts were dried at 4 °C using a CentriVap concentrator, derivatized by methoximation and silylation, and analyzed by gas chromatography–mass spectrometry (GC–MS) on an Agilent 7890A GC coupled to a 5975C mass spectrometer. Metabolites were identified based on characteristic fragmentation patterns and retention times, and isotopic enrichment was determined by integrating diagnostic ion fragments according to published references^29,30^. Natural abundance of stable isotopes was corrected for using the R-based tool IsoCorrectoR.

### Statistics

Data are presented as mean ± SEM, unless otherwise indicated. Statistical analyses were performed using GraphPad Prism (version 11). Normality was assessed using the D’Agostino– Pearson omnibus test. For comparisons between two groups, unpaired two-tailed Student’s t-tests were used for normally distributed data, and Mann–Whitney tests were applied for non-normally distributed data. For comparisons involving more than two groups, one-way or two-way ANOVA was performed for normally distributed data, followed by Tukey’s multiple-comparison post hoc tests as appropriate. When normality was not met, Kruskal–Wallis tests with Dunn’s post hoc correction were applied. The number of independent biological replicates (n) is indicated in the figure legends. P values < 0.05 were considered statistically significant. For proteomic datasets, pairwise or multiple testing correction was performed using false discovery rate (FDR) adjustment as specified in the corresponding Methods section.

## Acknowledgements

We thank the technical assistant staff from the Stem Cell Unit (UMG), especially Yvonne Wedekind and Yvonne Hintz; Behnoush Parviz, Antje Weber, Stefanie Wolfram and Henrike Thomas from the Experimental and Translational Cardiology Institute (JLU Giessen), for superb technical support. Lisa Neuenroth from the Proteomics Core Unit (UMG) for her proteomics assistance.

## Author contribution

M.P-G designed the study. M.P-G, L.C and E.P-C and designed the experiments. K.a.d.B, J.F, T.S, F.L, H.A, J.P, C.L, A.C-O, L.C and M.P-G performed the experiments and analyzed the data. S.J, V.R.T, S.T.S, L.C.Z provided technical support and conceptual advice. M.P-G, L.C. and E.P-C wrote and edited the manuscript.

