## Extended data for "AMPK reinforces mitochondrial metabolism and suppresses pathological remodeling in Complex V–deficient cardiomyocytes"

**Esteban Palacios-Contreras**

**Karen an der Brügge**

**Jakob Fell**

**Till Stephan**

**Hugo Amedei**

**Christof Lenz**

**Jelena Pesek**

**R. Verena Taudte**

**Felix Lange**

**Stefan Jakobs**

**Samuel Sossalla**

**Laura C. Zelarayán**

**Alfredo Cabrera-Orefice**

**Lukas Cyganek**

**Mario G. Pavez-Giani**

### Extended material:

### Material and Methods.

#### Generation of *TMEM70*-deficient iPSC

The *wild-type* iPSC line UMGi014-C clone 14, designated $TMEM70^{WT/WT}$, was previously characterized and derived from dermal fibroblasts of a healthy male donor via integration-free Sendai virus transduction[1]. Genome editing in early passage iPSCs was performed using ribonucleoprotein-based CRISPR-Cas9 with crRNA/tracrRNA and Hifi SpCas9. To form the ribonucleoprotein complex, Alt-R CRISPR-Cas9 crRNA [300 pmol ], and Alt-R CRISPR-Cas9 tracrRNA [300 pmol] were pre-assembled with Alt-R Hifi SpCas9 Nuclease 3NLS [122 pmol] (all IDT DNA Technologies). Nucleofection was performed with $2 \times 10^6$ iPSCs using the 4D Amaxa Nucleofector system (Lonza; program CA-137) and the P3 Primary Cell 4D-Nucleofector X Kit (Lonza) according to manufacturer's instructions. Following nucleofection, iPSCs were replated into a Matrigel-coated (growth factor reduced, BD Biosciences) 6-well plate containing StemFlex medium (Thermo Fisher Scientific) supplemented with thiazovivin [2 µM] (Merck) and 100 U/ml penicillin and 100 µg/ml streptomycin (Thermo Fisher Scientific). After 3 days, transfected iPSCs were singularized using the CellenOne single cell dispenser (Cellenion/Scienion) in StemFlex medium on Matrigel-coated 96-well plates. Successful genome editing was identified by Sanger sequencing. For $TMEM70^{KO/KO}$ generation, WT iPSCs were genome-edited by CRISPR-Cas9, targeting *TMEM70* exon 1 using guide RNA target sequence 5'- TCGAACTGCCTCTCTGCGGA**AGG**-3' (PAM in bold), and the CRISPR-edited isogenic iPSC lines UMGi014-C-28 clone 17 was established. For $TMEM70^{SNP/SNP}$ insertion of c.317-2A>G TMEM70 biallelic, WT iPSC line UMGi014-C clone 14 was genome-edited by CRISPR-Cas9, targeting *TMEM70* intron 2 using guide RNA target sequence 5′- AGAAACATTTCACACCTAAA**TGG**-3' (PAM in bold), in combination with a single-stranded oligonucleotide for homology-directed repair, and the CRISPR-edited isogenic iPSC line was established as UMGi014-C-29 clone 3.

Generated iPSC lines were maintained on Matrigel-coated plates, passaged every 4-6 days with Versene solution (Thermo Fisher Scientific) and cultured in StemMACS iPS-Brew XF medium (Miltenyi Biotec) supplemented with 2 µM thiazovivin on the first day after passaging with daily medium changes. Cultures were maintained in a humidified atmosphere (95% air, 5% CO2) at 37°C. Pluripotency analysis via immunocytochemistry and flow cytometry was performed as previously described[1]. For molecular karyotyping, genomic DNA from iPSC clones was sent for genome-wide analysis using Illumina BeadArray (Life&Brain, Germany). Digital karyotypes were analyzed in GenomeStudio v2.0 software (Illumina) and visualized using Circa (OMGenomics Labs). Copy number events were reported if larger than 3.5×105 bps and 1×106 bps for loss of heterozygosity.

**iPSC-derived cardiomyocyte differentiation**

Functional ventricular iPSC-derived cardiomyocytes (iPSC-CMs) were generated from human iPSC lines, including included $TMEM70^{WT/WT}$, mono- ($TMEM70^{KO/WT}$ and $TMEM70^{SNP/WT}$) and biallelic loss of *TMEM70* ($TMEM70^{KO/KO}$ and $TMEM70^{SNP/SNP}$). The methodology involved WNT signaling modulation and a subsequent metabolic selection process, adapted from prior methodology[2]. Briefly, iPSC-CM differentiation was initiated at a cellular confluence of 80-90% in Matrigel-coated plates with cardio differentiation medium composed of RPMI 1640 with GlutaMAX and HEPES (Thermo Fisher Scientific), 0.5 mg/ml human recombinant albumin (Merck) and 0.2 mg/ml L-ascorbic acid 2-phosphate (Merck), and sequential treatment with 4 µM CHIR99021 (Merck) for 48 h and 5 µM IWP2 (Merck) for further 48 h. On day 8, medium was changed to cardio culture medium composed of RPMI 1640 with GlutaMAX and HEPES, and supplemented B27 (Thermo Fisher Scientific), with medium changed every two days. Differentiated iPSC-CMs cultures around day 11 were metabolically selected using cardio selection medium composed of RPMI 1640 without glucose (Thermo Fisher Scientific), 0.5 mg/ml human recombinant albumin, 0.2 mg/ml L-ascorbic acid 2-phosphate and 4 mM lactate (Merck) for 7 days. After 48 h of recovery in cardio culture medium, iPSC-CM were digested with 0.25% Trypsin/EDTA (Thermo Fisher Scientific) and replated at lower density (1:2-1:3) in Matrigel-coated plates. iPSC-CMs were maintained in feeder-free and serum-free culture conditions until day 60 post-differentiation before being used for molecular and cellular experiments.

**SDS-PAGE and immunoblot**

Cell proteins were extracted from cultured either iPSC or iPSC-CMs using 1% of Sodium deoxycholate (SDC) pre-mixed with Pierce™ protease and phosphatase inhibitor mini tablets (Thermo Fischer Scientific) for at least 10 min. Protein containing supernatant was collected by centrifugation and stored at -80°C. Protein concentrations were determined using the Pierce BCA Protein Assay Kit (Thermo Fisher Scientific). Lysates were subjected to immunoblot analysis and to global proteome profiling (see below). For Immunoblot, samples were diluted to 1 µg/µl in sample buffer (final solution contained 10% glycerol, 50 mM DTT and bromophenol blue). Samples were heated to 95°C for 5 min. Proteins were separated in either Mini-PROTEAN TGX™ or Criterion TGX™ Precast Gels (Bio Rad) using 1× tris/glycine/SDS buffer (Bio-Rad). Next, proteins were transferred to nitrocellulose membranes using Trans-Blot

Turbo RTA Transfer kit (Bio-Rad). Membranes were then blocked with a polyvinylpyrrolidone based solution (Carl-Roth) and incubated with primary antibodies at 4°C overnight and secondary HRP-labeled antibodies at room temperature for 1 h, before detecting with electrochemiluminescence. Total protein was used as loading control. Antibodies are listed in Supplementary Table 1.

**Global proteome profiling**

Two independent global proteome datasets were acquired in this study. The first dataset (library-based DIA) comprised fully differentiated iPSC-CMs (Fig. 2) and iPSC-CMs harvested on day 35 and day 65 of differentiation (Extended Data Fig. 5). The second dataset (library-free DIA) comprised treated iPSC-CMs treated either with vehicle or MK8722 (Fig. 4).

For *library-based DIA* (Fig. 2 and Extended data Fig. 5), iPSC-CMs pellets were lysed in a buffer containing 1% sodium deoxycholate (SDC) in [100 mM] (4-(2-hydroxyethyl)-1-piperazineethanesulfonic acid) (HEPES) assisted by sonication (Bioruptor pico). Protein extracts were cleaned and desalted using a magnetic bead-based SP3 protocol on amine-coated paramagnetic beads MagResyn Amine, Resyn (Biosciences)[3]. Following reduction, alkylation and tryptic digestion off the SP3 beads, peptides were dried in a Speedvac and kept at -20°C for further analysis. All samples were spiked with a synthetic peptide standard used for retention time alignment (iRT Standard).

Samples were redissolved in 20 µl loading buffer ([2%] acetonitrile, [0.1%] trifluoroacetic acid in water) spiked with a synthetic peptide standard used for retention time alignment (iRT Standard), and analyzed on a nanoflow chromatography system nanoRSLC (Thermo Fisher Scientific) hyphenated to a hybrid timed ion mobility-quadrupole-time-of-flight mass spectrometer (timsTOF Pro 2, Bruker). In brief, 400 ng equivalents of peptides were enriched on a reversed-phase C18 trapping column column [0.3 cm × 300 µm] (Thermo Fisher Scientific), and separated on a reversed-phase C18 column with an integrated CaptiveSpray Emitter [Aurora 3 series 25 cm × 75 µm] (IonOpticks) using a 100 min linear gradient of 5-34% acetonitrile/0.1% formic acid (v:v) at 200 nl $min^{-1}$, and a column temperature of 50ºC.

DIA analysis was performed in diaPASEF mode[4] using a 20x2 variable size window acquisition method from m/z 400 to 1,200 to include the 2+/3+/4+ population in the m/z–ion mobility plane. The collision energy was ramped linearly as a function of the mobility from 59 eV at 1/K0=1.6Vs cm−2 to 20 eV at 1/K0=0.6Vs cm−2. Two technical replicates per biological replicate were acquired.

Protein detection was performed in Spectronaut Software 18.5 (Biognosys) against an in-house annotated MS/MS spectral library for hiPSC-CMs generated against the against the UniProtKB

Homo sapiens reference proteome (revision 08-2023) augmented with an in-house protein contaminant database, using default parameters and at 1% FDR. For quantitation of DIA data, up to the 6 most abundant fragment ion traces per peptide, and up to the 10 most abundant peptides per protein were integrated and summed up to provide protein area values. Mass and retention time calibration and the corresponding extraction tolerances, were dynamically determined. Both identification and quantification results were trimmed to a False Discovery Rate (FDR) of 1% using a forward-and-reverse decoy database strategy.

For *library-free DIA* (Fig.4), cells were pelleted in 1.5 ml Eppendorf tubes by centrifugation, snap-frozen in liquid nitrogen, and stored at -80 °C until proteome analysis. For protein extraction, cells were thawed on ice and resuspended in the SDC-containing lysis buffer described above, supplemented with 0.125 U/µl benzonase. After thorough vortexing and 15 min incubation at room temperature, small aliquots were taken for protein concentration determination by Pierce BCA assay.

Aliquots containing 100 µg of protein were used for protein digestion and further peptide cleanup using the iST kit (PreOmics) according to the manufacturer's instructions with slight modifications. Briefly, protein pellets were solubilized in 50 µl of LYSE buffer by thorough vortexing and boiling at 95 °C for 10 min with shaking at 1000 rpm. This step included reduction and alkylation of cysteines. Proteins were digested with Trypsin/LysC overnight at 37 °C. Peptides were cleaned using iST cartridges and eluted into a 96-well microtiter plate. After complete drying at 45 °C using a Concentrator Plus (Eppendorf), peptides were resuspended in 20 µl of LC-LOAD buffer by vortexing.

Peptides were analyzed by LC-MS/MS using an UltiMate 3000 RSLCnano system (Thermo Fisher Scientific) coupled to an Orbitrap Eclipse Tribrid mass spectrometer. For each sample, 5 µl of peptides were loaded onto a PepMap Neo Trap column (Thermo Fisher Scientific) for concentration and desalting, followed by separation on an Extend column (1.9 µm C18, 25 cm length × 150 µm inner diameter; ProtumLink GmbH, Germany) using an Easy-Spray adapter. The analytical column was maintained at 60 °C. Peptide separation was achieved with a linear gradient of solvent A (0.15% formic acid in HPLC-grade water) and solvent B (0.15% formic acid in HPLC-grade acetonitrile) at a flow rate of 0.5 µl/min given at min (%B): 0-5 (const. 3%), 5-10 (3–10%), 10-145 (10-25%), 145-158 (25-35%), 158-165 (35-45%) and 165-168 (45-85%). The column was rinsed with 85% B for 5 min at 0.8 µl/min and equilibrated for 3 and 4 min with 3% B at 0.8 and 0.5 µl/min, respectively. The total time of each run was 180 min. MS analysis was performed in positive ion mode with electrospray ionization (2.2 kV) and source temperature set to 275 °C. Data were acquired in data-independent acquisition (DIA) mode using HCD fragmentation. MS1 spectra were recorded over a 350–1650 m/z range at 120K resolution in the Orbitrap, with an AGC target of $1.2 \times 10^6$ and a maximum injection time of 20 ms. RF Lens was set to 40% and MS1 spectra were acquired in profile mode. DIA settings were as follows: precursor mass scan range: 375-1200 m/z; isolation mode: quadrupole; isolation window (m/z): 10; scan range mode: define first mass; first mass (m/z): 200; loop control: 2; loop time (s): 3; loop count: 20; HCD collision energy/energies (%): 25.5, 27, 30; Orbitrap resolution: 30K; automatic gain control value set to 1.5E6 ions; maximum injection time of 55 ms. Raw data were analyzed in a single run using DIA-NN (v2.2.0) in library-free mode[5]. Trypsin was selected as the protease with one missed cleavage allowed. Cysteine carbamidomethylation and N-terminal methionine excision were enabled. The options for unrelated runs, match-between-runs, and protein inference were activated. The quantification strategy was set to Legacy (direct) and cross-run normalization was disabled. The false discovery rate (FDR) was set to 1% (default). Spectral library generation used the UniProt *H. sapiens* reference proteome (June 2025), including a default list of contaminants. All other parameters were kept at default values. The resulting report.pg matrix output file was filtered to exclude contaminants.

For both datasets, differentially expressed proteins were identified using statistical thresholds adjusted for pairwise or multiple testing. Gene ontology pathway enrichment analysis of differentially abundant proteins ($p \leq 0.05$) was performed using Enrichr (Ma'ayan Laboratory). Downstream analysis and visualization were carried out in Microsoft Excel 365 and an in-house R/Shiny tool. When needed, data visualization was developed with the assistance of Claude (Anthropic); all analytical approaches, outputs, and interpretations were independently reviewed and validated by the authors.

### Blue Native PAGE and complexome profiling

Whole cells and mitochondria isolated by differential centrifugation were solubilized under native conditions and separated by Blue Native PAGE according to Wittig et al. 2006[6]. Whole cells were solubilized in a 1 % NP40-containing buffer, while mitochondria were solubilized in a digitonin-containing buffer at a digitonin-to-protein ratio of 6 g/g; both were separated on 3–12% acrylamide gradient gels. Complex V assembly was analyzed by immunoblotting after Blue Native PAGE using an antibody against the ATP synthase subunit ATP5F1A.

For complexome profiling, gel lanes were sliced into 35 and 40 fractions for whole-cell and mitochondrial samples, respectively. Following electrophoresis, gels were fixed overnight in 50% methanol, 10% acetic acid, 10 mM ammonium acetate, stained with 0.025% Coomassie

Brilliant Blue G-250 (SERVA) in 10 % acetic acid, and subsequently destained in 10% acetic acid. Gels were stored in deionized water to allow complete re-swelling prior to imaging.

Digitized full-size gel images were used as templates to guide systematic slicing. Individual lanes were cropped and sectioned from bottom to top into equal fractions as indicated above. Gel slices were diced and transferred into 96-well filter plates (Merck Millipore). Coomassie dye was removed by repeated washing with 50% methanol in 50 mM ammonium bicarbonate, including intermediate centrifugation steps to remove excess liquid.

Proteins were reduced with 10 mM TCEP and alkylated with 40 mM chloroacetamide prior to in-gel digestion. After air-drying, gel pieces were rehydrated with 40 µL of digestion solution (2.5 ng Trypsin/µL, 50 mM ammonium bicarbonate, 1 mM $CaCl_2$), incubated for 15 min at 4 °C to allow enzyme uptake, supplemented with 60 µL of 50 mM ammonium bicarbonate to cover the gel pieces and digested overnight at 37 °C. Peptides were recovered by centrifugation followed by a second extraction with 30 % acetonitrile, 3 % formic acid. Combined eluates were vacuum-dried and reconstituted in 2% acetonitrile and 0.5% formic acid prior to LC–MS/MS analysis.

Peptide fractions were analyzed using an Ultimate 3000 UHPLC system coupled to an Orbitrap Eclipse Tribrid mass spectrometer (Thermo Fisher Scientific). Prior to analytical separation, peptides were concentrated and desalted on a PepMap™ Neo Trap Cartridge (Thermo Fisher Scientific). Chromatographic separation was performed using an Extend column (1.9 µm C18, 25 cm length × 150 µm inner diameter; ProtumLink GmbH, Jena, Germany). Peptides were eluted at 500 nL/min using a 28 min gradient with solvent A (0.1% formic acid in water) and solvent B (0.1% formic acid in acetonitrile) as follows: loading 0–5 min, 5 %B; separation: 5–7.5 min, 5–10 %B, 7.5–30 min, 10–35 %B, and high-organic elution 30–33 min, 35–90 %B. The column was subsequently washed at 800 nL/min for 5 min (90 %B), returned to 5 %B within 1 min, and re-equilibrated for 5 min at 500 nL/min. Total run time per fraction was 45 min.

Mass spectrometry data were acquired in positive ion mode and data-dependent acquisition (DDA) mode following a Top20 method, in which full MS1 spectra were acquired in the Orbitrap at a resolution of 75,000 (at *m/z* 200) over a range of *m/z* 375–1500, with an AGC target of $3 \times 10^6$, automatic maximum injection time, and the RF lens set to 30%. Spectra were recorded in profile mode. The 20 most intense precursor ions were isolated in the quadrupole (isolation window = 1.6 *m/z*) and subjected to collision-induced dissociation (CID) with a normalized collision energy of 35%, followed by fragment ion analysis in the ion trap in rapid scan mode, with a normalized AGC target of 250% and dynamic maximum injection time.

Raw data from all gel fractions were processed using MaxQuant (v2.7.1.0) against the UniProt reference proteome of *H. sapiens* (June 2025). Carbamidomethylation of cysteine residues was specified as a fixed modification, while methionine oxidation, deamidation (NQ), and protein N-terminal acetylation were included as variable modifications. Match between runs (1 min) was enabled, while all other parameters were kept as default. Protein identifications were filtered at a false discovery rate of 1 %. Relative protein abundance across gel slices was estimated using intensity-based absolute quantification (iBAQ) values. Migration profiles were generated for each protein group and normalized to the maximal signal intensity observed across all fractions. Hierarchical clustering based on Pearson correlation distance with average linkage was applied to group proteins according to their migration behaviour. Apparent molecular mass calibration of BN-gels was performed using reference membrane and soluble protein complexes. To account for loading differences and MS sensitivity, complexome datasets were scaled using the summed iBAQ values of all the identified proteins annotated in MitoCarta 3.0[7] excluding the subunits from the ATP synthase complex. For visualization, the maximal value of each protein group across gel slices was set to 1 and normalized accordingly. Data downstream analysis and visualization was done manually in Microsoft Excel 365.

**Mitochondrial respiration assay**

Oxygen consumption rate (OCR) was measured using a Seahorse XFe24 Analyzer (Agilent Technologies). $1 \times 10^5$ iPSC-CMs were seeded per well onto Matrigel-coated XFe24 plates seven days prior to the assay. Cartridge sensors were equilibrated in calibrant assay medium 24 h prior to measurement. Basal respiration was recorded, followed by sequential injection of oligomycin (ATP synthase inhibitor; [1µM]), FCCP (uncoupler; [1µM]), and rotenone/antimycin A (complex I/III inhibitors; [0.5 and 0.5µM, respectively]). ATP-linked respiration was calculated as the difference between basal OCR and oligomycin-treated OCR. Maximal respiration was defined as FCCP-stimulated OCR. Spare respiratory capacity was calculated as maximal minus basal respiration. Proton leak was defined as OCR remaining after oligomycin treatment minus non-mitochondrial respiration. OCR values were normalized to protein concentration using BCA methods. Normalized values ere analysis using Wave Pro Software (Agilent Technoligies).

**Mitochondrial potential assay**

Mitochondrial membrane potential (Δψm) was assessed using tetramethylrhodamine methyl ester (TMRM) (Invitrogen™) and MitoTracker Green (MTG) (ThermoFisher) for mitochondrial mass normalization. 1.5 x $10^4$ iPSCs or 1 x $10^5$ iPSC-CMs were seeded per well onto Matrigel-coated 24-well plates seven days prior to the assay iPSC-derivatives were treated with TMRM [20 nM] (non-quenching mode) and MTG [100 nM] for 20 min. After incubation competition, cells were washed three times with PBS and incubated with RPMI media without phenol red. Cells were visualized using Incucyte® automated-imaging system. Mean TMRM (red channel) fluorescence intensity per cell was quantified and normalized to MTG (green channel) signal. For depolarization controls, FCCP [5 μM ] was applied acutely.

**STED microscopy**

Cells were seeded onto 18 mm #1.5H coverslips (Marienfeld Superior, Lauda-Königshofen, Germany) pre-coated with Matrigel. Cells were fixed in pre-warmed (37 °C) 8% formaldehyde (Merck Millipore, Darmstadt, Germany; Cat. No. 30525-89-4) in PBS (pH 7.4) for 5 min, washed with PBS, permeabilized with 0.5% Triton X-100 in PBS for 5 min, and blocked with 5% bovine serum albumin (BSA; Merck Millipore, Darmstadt, Germany) in PBS for 20 min.

Samples were incubated for 1.5 h at room temperature with 5% BSA containing Phalloidin–Oregon Green 488 (Invitrogen, Waltham, MA, USA; O7466; 1:100) and primary antibodies against MIC60 (Abcam, Cambridge, UK; ab137057; 1:100) and ATP5F1B/ATP5B (Abcam; ab5432; 1:200), followed by five PBS washes. Primary antibodies were detected using goat anti-rabbit secondary antibodies (Jackson ImmunoResearch Laboratories, West Grove, PA, USA; 111-005-144) custom-labeled with Abberior STAR RED (Abberior, Göttingen, Germany) and goat anti-mouse Alexa Fluor 594 secondary antibodies (Life Technologies, Waltham, MA, USA; A11005), both diluted 1:100 in PBS and incubated for 1 h at room temperature. After five PBS washes, samples were rinsed with distilled water and mounted in Mowiol supplemented with DABCO.

STED images were acquired on a Quad-Scanning STED microscope (Abberior Instruments, Göttingen, Germany) equipped with a UPlanSApo 100×/1.40 Oil objective (Olympus, Tokyo, Japan) and controlled using Imspector 16.1-win64-AIFpgaV3 software. Oregon Green, Alexa Fluor 594, and Abberior STAR RED were excited at 485, 561, and 640 nm, respectively. STED imaging was performed using pulsed depletion at 775 nm in time-gated mode. Fluorescence was detected with a 0.75 ns delay after the STED pulse over an 8 ns detection window. Pixel size was 18–20 nm, the pinhole was set to 0.8–1 Airy units (AU), the pixel dwell time to 10–15 μs, and line accumulation to 3–5 lines. Raw image data were processed using the low-pass filter implemented in Imspector, and mitochondrial morphology metrics were quantified in ImageJ.

**Electron microscopy**

iPSC-CMs were immobilized with a solution containing 2.5% glutaraldehyde in 0.1M cacodylic [0.1M] buffer (pH 7.4) (Science Services) for one hour at room temperature. The fixation was completed over night at 4°C. The samples were washed with [0.1M] cacodylic buffer (pH 7.4) at room temperature for 5 min each using a rocking platform. Subsequently, the samples were incubated in a solution containing 1.5% potassium-hexacyanoferrate (Merck KGaA) and 1% OSO4 (Science Services) in cacodylic [0.1M] buffer (pH 7.4) at room temperature for one hour. Without washing, the samples were further incubated in a solution of 1% $OsO_4$ in 0.1 M cacodylic buffer (pH 7.4) at room temperature for one hour. The samples were then washed three times with ddH2O for 10 min each before incubation with a solution of 1% uranyl acetate (Science Services) in ddH2O at room temperature for 30 min in the dark. The samples were washed three times with ddH2O for 5 min each and subsequently dehydrated with a graded ethanol series. The final dehydration was performed in propylene oxide (Merck KGaA) twice for 10 minutes each. Dehydrated samples were infiltrated with EMbed812 (Science Services) diluted 1:1 with propylene oxide for one hour at room temperature on a rocking platform. The samples were then placed in pure EMbed812 resin, placed in an evacuated desiccator and incubated for one hour at room temperature on a rocking platform. The resin was exchanged for pure resin, placed in an evacuated desiccator and incubated overnight at room temperature on a rocking platform. Finally, samples were covered with beem capsules in fresh resin and cured at 60°C for 48 hrs. Sections of 70nm thickness were obtained from the samples and collected on Formvar/carbon coated grids and examined on a Phillips CM120 transmission electron microscope (FEI Company) operated at 120kV and equipped with a TVIPS TemCam F416 camera (TVIPS).

**NanoString**

For gene panel analysis, a customized nCounter Elements TagSet panel designed by NanoString Technologies was utilized. 200 ng of RNA per sample were hybridized to target-specific capture and reporter probes at 67°C for 16-20 h according to the manufacturer’s instructions. Samples were then loaded into the NanoString cartridge, and the nCounter gene expression assay was initiated. Raw reads were analyzed with nSolverTM Data Analysis Software. For background subtraction the geometric means of negative controls was used. RNA counts were normalized to four housekeeping genes (TBP, HPRT1, POL2RA, and GAPDH). TagSets are listed in Supplementary Table 2.

**Contractility measurements in spontaneously beating iPSC-CMs**

Spontaneously beating cardiomyocytes were recorded using phase-contrast video microscopy (IonOptix). Contractile parameters, including beating frequency, contraction amplitude, and time to 50% peak contraction, were analyzed using motion-tracking software. At least two independent differentiations were analyzed per condition.

**Cytotoxicity assay**

To analyze cell viability, $1 \times 10^5$ iPSC-CMs were seeded onto a 24-well plate. After 7 days of recovery, cells were treated with different concentrations of MK8722 or the vehicle control (DMSO 1:1,000) for an additional 7 days, administered during media change every other day. Before imaging acquisition, Calcein AM Viability Dye [1 µM] (Thermo Fisher Scientific) in RPMI medium without phenol red was administrated to the cultures for 30 min at 37°C. Subsequently, cells were washed twice with with PBS, incubated with propidium iodide [500 nM] (Merck) in RPMI medium without phenol red, and analyzed using the IncuCyte S3 System to discriminate between live and dead cells.

**Carbon tracing and metabolomic analysis**

For carbon tracing experiments [U-$^{13}C_6$], iPSC-derived cardiomyocytes were incubated in either glucose-free RPMI supplemented with B27, 2 mM L-glutamine, and [U-$^{13}C_6$]glucose [10 mM] or RPMI supplemented with B27, 2 mM L-glutamine, carnitine [0.5mM], and [U-$^{13}C_6$]Palmitic-acid [100 µM] conjugated with BSA [1:3 ratio]. MK8722 [1 µM] was maintained throughout the labeling period. iPSC-CMs were labeled for the specified durations under standard culture conditions. Metabolism was quenched by rapid aspiration of medium followed by immediate addition of ice-cold 80% (v/v) methanol. Cells were incubated on dry ice, scraped into extraction solvent and centrifuged at 16,000g for 10 min at 4 °C. The extracts were dried at 4 °C using a CentriVap concentrator (Labconco). Dried metabolites were reconstituted in 20 µL of methoxyamine hydrochloride solution (2 % in pyridine) and incubated for 1.5 hours at 40 °C. Derivatization was then carried out with 20 µL of MTBSTFA for 3 hours at 50 °C. Gas chromatography–mass spectrometry (GC–MS) analysis was performed on an Agilent 7890A gas chromatograph coupled to a 5975C mass spectrometer. A 1 µL aliquot of each sample was injected in splitless mode onto an HP-5MS column (30 m × 0.25 mm, 0.25 µm film thickness) with helium as the carrier gas (1 mL $min^{-1}$). The inlet and transfer line temperatures were maintained at 270 °C and 280 °C, respectively. The oven temperature was programmed to increase from 100 °C (2 min hold) to 300 °C (1 min hold) at a rate of 3.5 °C $min^{-1}$. The MS source and quadrupole were held at 230 °C and 150 °C, respectively, and electron impact ionization was applied at 70 eV. Spectra were acquired in full-scan mode, and data were processed using MassHunter Quantitative Analysis (version B.05.00). Metabolites were identified based on their characteristic fragmentation patterns and retention times, and isotopic enrichment was determined by integrating diagnostic ion fragments according to published references [8,9].

To correct for the natural abundance of stable isotopes the R-based tool IsoCorrectoR was used.

**Statistics**

Data are presented as mean ± Standard error of the mean (SEM), unless otherwise indicated. Statistical analyses were performed using GraphPad Prism (version 11). Normality was assessed using the D'Agostino–Pearson omnibus test. For comparisons between two groups, unpaired two-tailed Student's t-tests were used for normally distributed data, and Mann–Whitney tests were applied for non-normally distributed data. For comparisons involving more than two groups, one-way or two-way ANOVA was performed for normally distributed data, followed by Tukey's multiple-comparisons post hoc tests as appropriate. When normality was not met, Kruskal–Wallis tests with Dunn's post hoc correction were applied. The number of independent biological replicates (n) is indicated in the figure legends. P values $< 0.05$ were considered statistically significant. For proteomic datasets, pairwise or multiple testing correction was performed using false discovery rate (FDR) adjustment as specified in the corresponding Methods section.

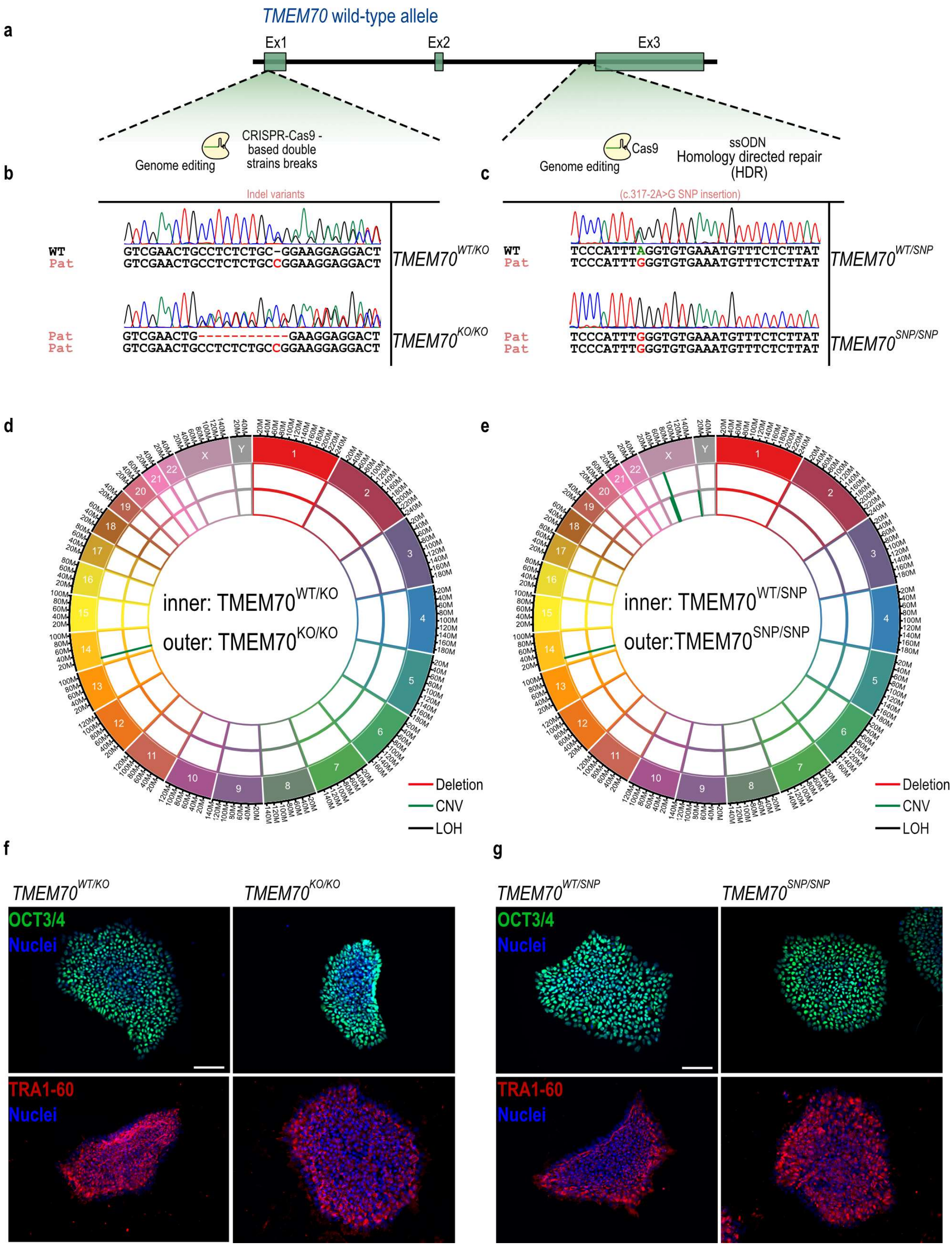


**Extended Data Figure 1. Generation and characterization of TMEM70-deficient iPSC lines.**

(a) Schematic of the TMEM70 *wild-type* allele (exons 1–3, Ex1–Ex3) and CRISPR-Cas9 genome-editing strategy. Frameshift indel variants were introduced at exon 1 via Cas9-induced double-strand breaks. The patient-mimicking splice-site variant (c.317-2A>G) was introduced at intron 2 via Cas9 in combination with a single-stranded oligodeoxynucleotide (ssODN) donor for homology-directed repair (HDR).

(b) Sanger sequencing chromatograms confirming indel editing at exon 1. Top: heterozygous single-nucleotide insertion generating TMEM70$^{WT/KO}$. Bottom: biallelic editing (single-nucleotide insertion on one allele, multi-nucleotide deletion on the second) generating TMEM70$^{KO/KO}$. WT, *wild-type* reference sequence; Pat, pathological/edited allele sequence; inserted or altered bases highlighted in red.

(c) Sanger sequencing chromatograms confirming introduction of the c.317-2A>G splice-site variant in intron 2. Top: heterozygous A>G substitution generating TMEM70$^{WT/SNP}$. Bottom: homozygous G/G genotype generating TMEM70$^{SNP/SNP}$. Substituted base highlighted in red.

(d, e) Genome-wide copy-number/genomic integrity analysis confirming normal karyotype and absence of large-scale copy-number variants in TMEM70$^{WT/KO}$ and TMEM70$^{KO/KO}$ lines (d) and TMEM70$^{WT/SNP}$ and TMEM70$^{SNP/SNP}$ lines (e) following CRISPR-Cas9 editing. Chromosomes 1–22, X and Y are shown; the scale indicates a genomic position in megabases (M). CNV, copy number variants; LOH, loss of heterozygosity.

(f, g) Immunofluorescence staining for pluripotency markers OCT3/4 (green) and TRA-1-60 (red), with nuclear counterstain with DAPI (blue), confirming maintained pluripotency in TMEM70$^{WT/KO}$ and TMEM70$^{KO/KO}$ (f) and TMEM70$^{WT/SNP}$ and TMEM70$^{SNP/SNP}$ (g) iPSC colonies following genome editing. Scale bar, 100 μm.

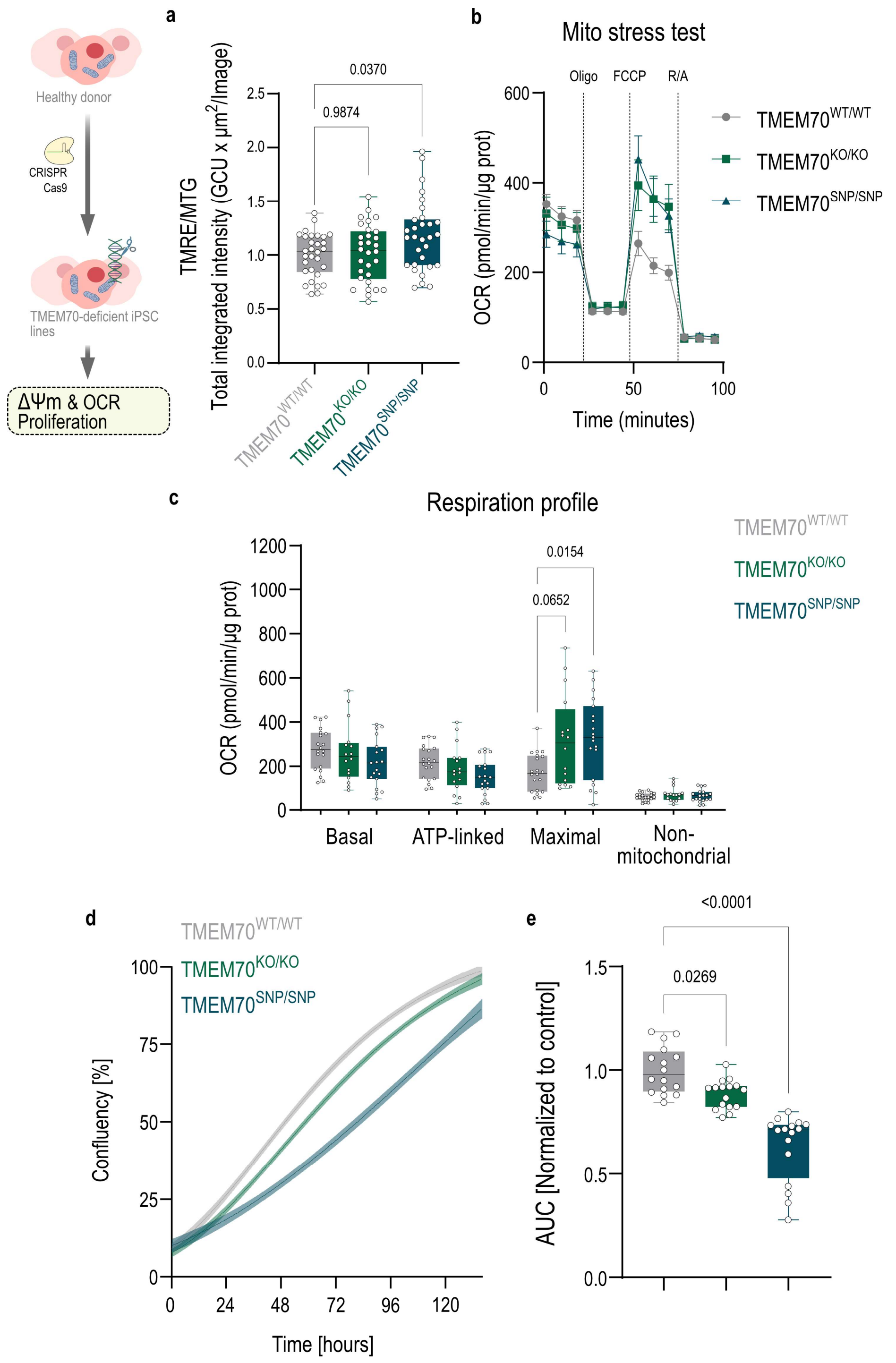


**Extended Data Figure 2. TMEM70-deficient iPSC lines present a mild, largely compensated phenotype at the pluripotent stem cell stage.**

(a) Mitochondrial membrane potential (Δψm), quantified as TMRE/MTG total integrated intensity, in TMEM70$^{WT/WT}$, TMEM70$^{KO/KO}$, and TMEM70$^{SNP/SNP}$ iPSCs. n=30 from five independent rounds. P values were determined by one-way ANOVA.

(b) Oxygen consumption rate (OCR) kinetics during mitochondrial stress test for $TMEM70^{WT/WT}$ $TMEM70^{KO/KO}$, and $TMEM70^{SNP/SNP}$ iPSCs with sequential addition of oligomycin, FCCP and rotenone/antimycin A.

(c) Quantification of basal, ATP-linked, maximal, and non-mitochondrial respiration in $TMEM70^{WT/WT}$, $TMEM70^{KO/KO}$, and $TMEM70^{SNP/SNP}$ iPSC lines normalized by total protein quantification. n=20, 16 and 19 respectively from three independent rounds. P values were determined by one-way ANOVA.

(d) Proliferative capacity of $TMEM70^{WT/WT}$, $TMEM70^{KO/KO}$, and $TMEM70^{SNP/SNP}$ iPSCs, measured as culture confluence (%) over seven days by live-cell imaging. Data shown as mean ± SEM.

(e) Quantification of (d) as area under the curve (AUC) of $TMEM70^{KO/KO}$ and $TMEM70^{SNP/SNP}$, relative to $TMEM70^{WT/WT}$ control. n=16 from four independent rounds; boxes show median and interquartile range with min–max whiskers. P values from one-way ANOVA with post hoc multiple comparisons.

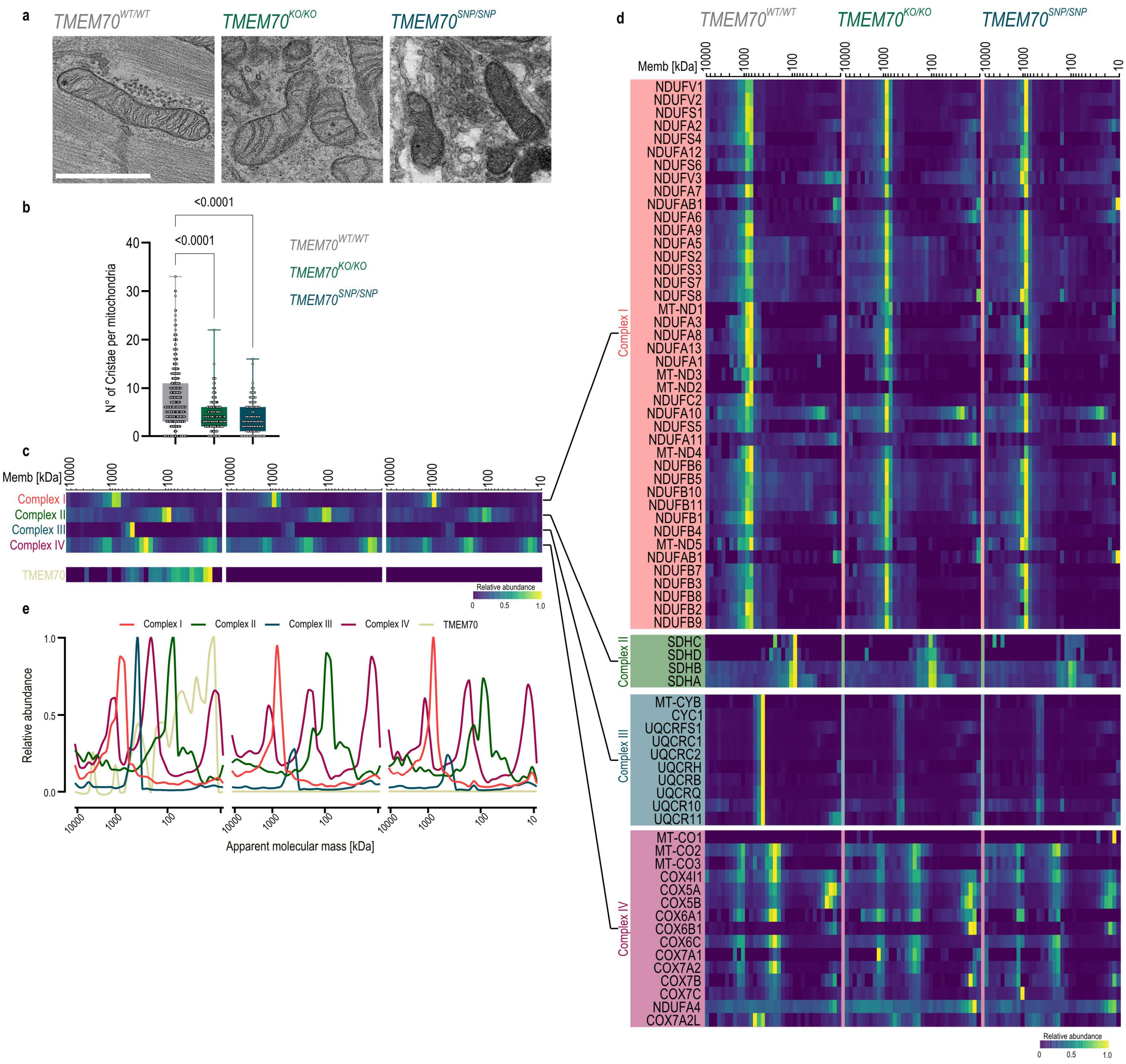


**Extended Data Figure 3. *TMEM70*-deficient cardiomyocytes exhibit disrupted mitochondrial cristae ultrastructure, and complexome profile reveals selective impairment of Complex V assembly.**

(a) Representative transmission electron microscopy images of mitochondria in TMEM70$^{WT/WT}$, TMEM70$^{KO/KO}$, and TMEM70$^{SNP/SNP}$ cardiomyocytes, showing disorganized cristae architecture in TMEM70-deficient cells compared to well-organized parallel cristae in wild-type controls. Scale bar, 100 nm.

(b) Quantification of the number of cristae per mitochondrion. Each data point corresponds to a single mitochondrion observed in TMEM70$^{WT/WT}$ (n=422), TMEM70$^{KO/KO}$ (n=183), and TMEM70$^{SNP/SNP}$ (n=104). Analysis was performed from two independent differentiations. P values were determined by Kruskal-Wallis test.

(c) Complexome profiling of complexes I-IV in *TMEM70*-deficient iPSC-CMs. Heatmaps show the relative abundance and distribution of ETC complexes across assembly states.

(d) Heatmap of complexome profiling for individual subunits of complexes I-IV across the three genotypes, corresponding to the complex-level summary in (c). Color scale indicates relative abundance (0–1).

(e) Migration profiles derived from (c), plotting relative abundance against apparent molecular mass for Complex I-IV in TMEM70-deficient iPSC-CMs compared with $TMEM70^{WT/WT}$.

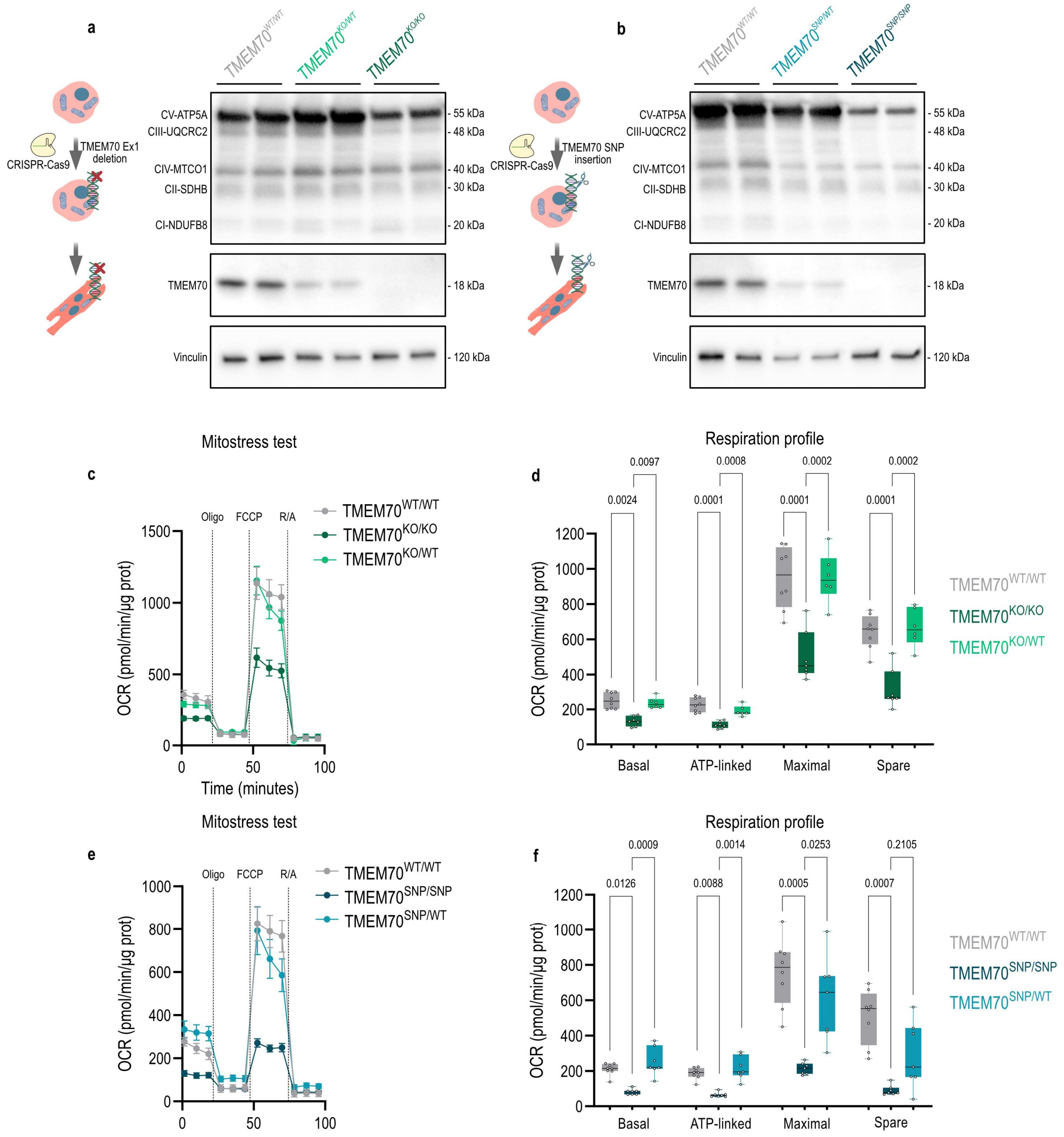


**Extended Data Figure 4. Monoallelic TMEM70 loss preserves Complex V protein levels and mitochondrial respiratory function.**

(a) Validation of TMEM70 expression in TMEM70$^{WT/WT}$, TMEM70$^{KO/WT}$, and TMEM70$^{KO/KO}$ at the iPSC-CMs. Immunoblot analysis was performed using specific antibodies against OXPHOS subunits of Complex I–V (top) and TMEM70 (middle). Vinculin served as loading control (bottom). Schematic (left) depicts CRISPR-Cas9-mediated deletion of TMEM70 exon 1.

(b) As in (a), for TMEM70$^{WT/WT}$, TMEM70$^{SNP/WT}$, and TMEM70$^{SNP/SNP}$ iPSC-CMs. Schematic (left) depicts CRISPR-Cas9-mediated insertion of the c.317-2A>G splice-site variant.

(c) Oxygen consumption rate (OCR) kinetics during mitochondrial stress test in TMEM70$^{WT/WT}$, TMEM70$^{KO/KO}$, and TMEM70$^{KO/WT}$ iPSC-CMs, with sequential addition of oligomycin, FCCP and rotenone/antimycin A.

(d) Quantification of basal, ATP-linked, maximal, and spare respiration from (c) normalized to total protein; n=6-9 from one differentiation. P values were determined by Kruskal-Wallis test.

(e) Oxygen consumption rate (OCR) kinetics during mitochondrial stress test in TMEM70$^{WT/WT}$, TMEM70$^{SNP/SNP}$, and TMEM70$^{SNP/WT}$ iPSC-CMs, with sequential injections as in (c).

(f) Quantification of basal, ATP-linked, maximal, and spare respiration from (e) normalized by total protein levels. n =7-8 from one differentiation; boxes show median and interquartile range with min–max whiskers. P values were determined by Kruskal-Wallis test.

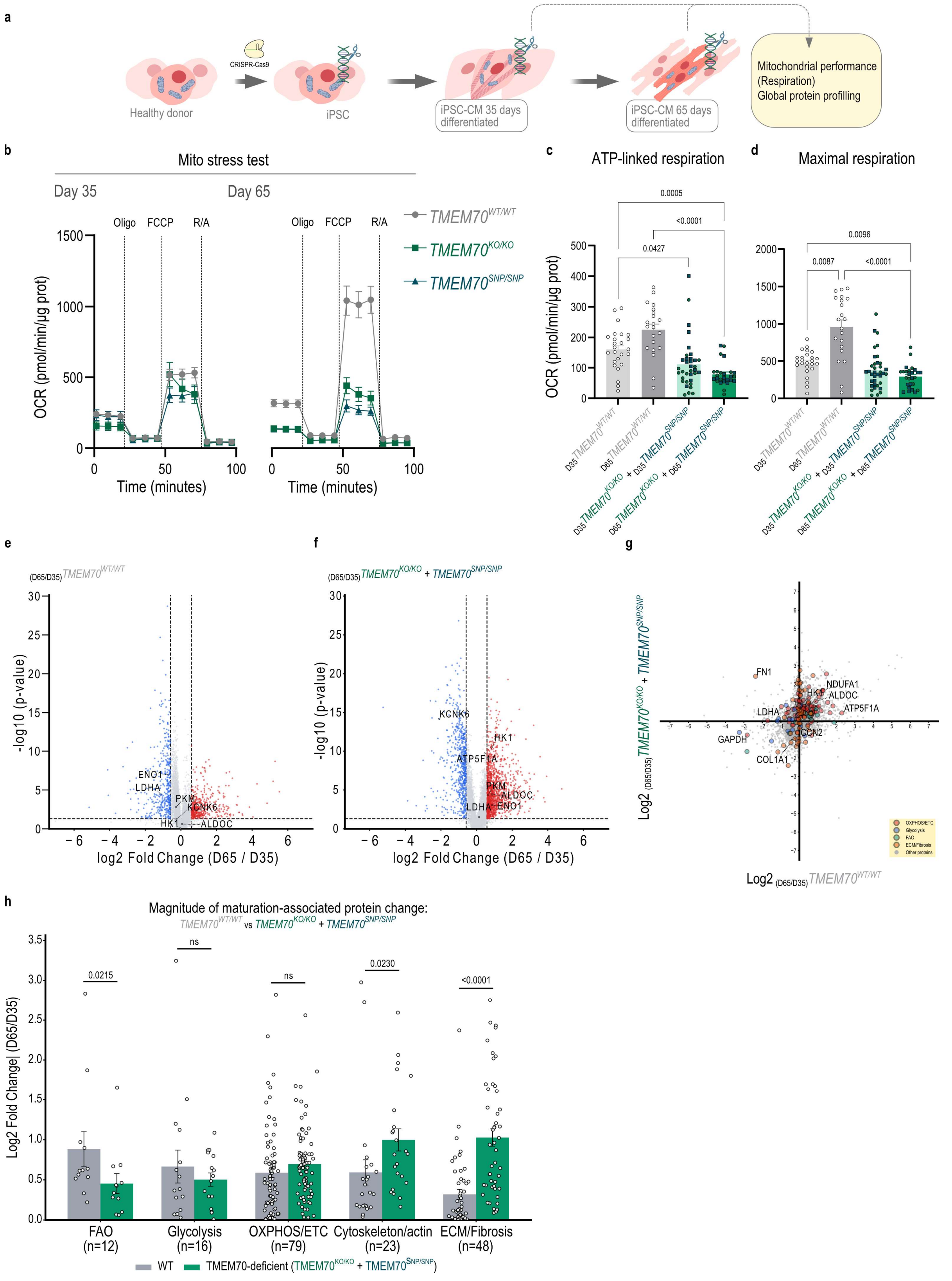


**Extended data Figure 5. *TMEM70* deficiency impairs maturation-associated respiratory capacity and drives a genotype-independent proteomic remodeling program**.

(a) Schematic overview of the two selected points during cardiomyocyte differentiation and the experimental workflow. *TMEM70*-deficient iPSC lines were differentiated into cardiomyocytes and analyzed at day 35 and day 65 of differentiation for oxygen consumption rate (OCR), and global proteome profiling.

(b) OCR kinetics during mitochondrial stress test at day 35 and day 65 of differentiation for $TMEM70^{WT/WT}$, $TMEM70^{KO/KO}$, and $TMEM70^{SNP/SNP}$ iPSC-CMs with sequential addition of oligomycin, FCCP and rotenone/antimycin A.

(c, d) Maximal respiration (c) and spare capacity normalized (d) at day 35 and day 65 for $TMEM70^{WT/WT}$, $TMEM70^{KO/KO}$, and $TMEM70^{SNP/SNP}$ iPSC-CMs. Total protein served as normalization. $TMEM70^{WT/WT}$; n=25 $_{(D35)}$, 21 $_{(D65)}$; $TMEM70^{KO/KO}$; n=18 $_{(D35)}$, 12 $_{(D65)}$; $TMEM70^{SNP/SNP}$: n=18 $_{(D35)}$, 14 $_{(D65)}$, pooled from three independent differentiations. P values were determined by Kruskal-Wallis test.

(e, f) Volcano plots of proteome-wide changes between day 65 and day 35 of differentiation in $TMEM70^{WT/WT}$ (e) and pooled Complex V-deficient ($TMEM70^{KO/KO}$ + $TMEM70^{SNP/SNP}$) (f) cardiomyocytes. Significance thresholds: |log2 fold change| > 0.585 (1.5-fold) and −log10(p-value) > 1.301 ($p < 0.05$), indicated by dashed lines. Red, significantly upregulated; blue, significantly downregulated; gray, not significant.

(g) Scatter plot comparing maturation-associated protein trajectories (log2 fold-change, day 65 vs. day 35) between $TMEM70^{WT/WT}$ (x-axis) and pooled Complex V-deficient ($TMEM70^{KO/KO}$ + $TMEM70^{SNP/SNP}$; y-axis) iPSC-CMs. Proteins are colored by pathway annotation: OXPHOS/electron transport chain (n=79), glycolysis (n=16), fatty-acid oxidation (FAO) (n=12), extracellular matrix (ECM)/fibrosis (n=48).

(h) Magnitude of maturation-associated protein change (mean |log2 fold change|, day 65 vs. day 35) for the pathway categories shown in (g), compared between $TMEM70^{WT/WT}$ and pooled TMEM70-deficient ($TMEM70^{KO/KO}$ + $TMEM70^{SNP/SNP}$) iPSC-CMs. Each point represents an individual protein; bars show mean ± SEM. P values from paired two-tailed t-tests comparing |log2FC| within each pathway between genotypes.

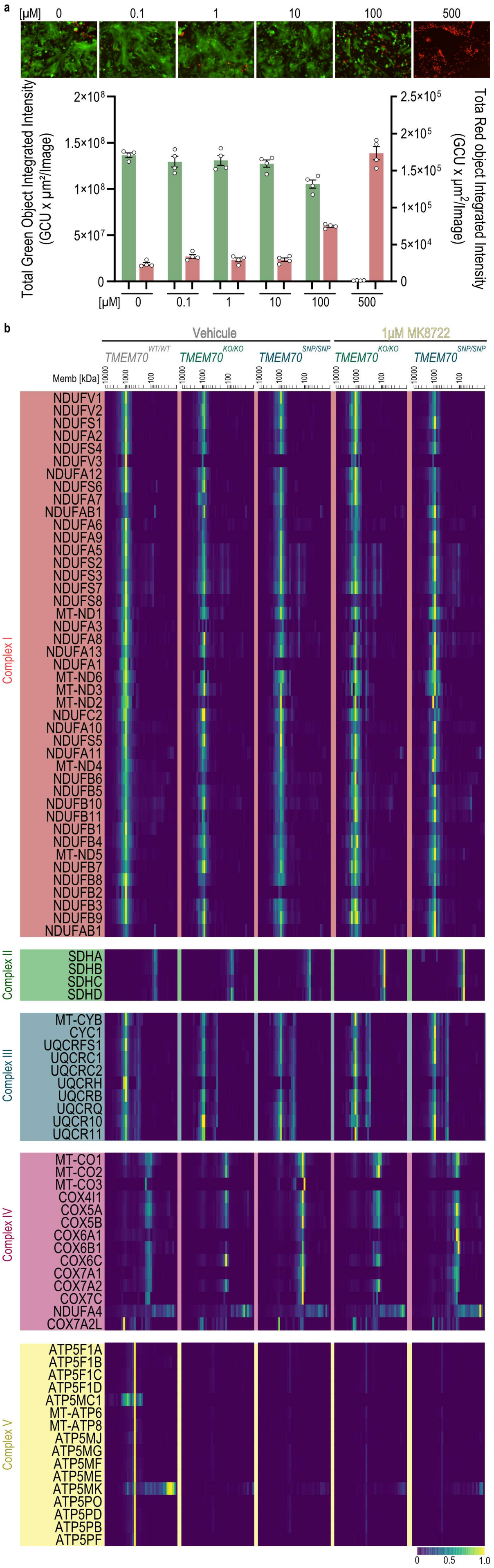

a
[µM] 0 0.1 1 10 100 500
Total Green Object Integrated Intensity (GCU x µm²/Image)
2×10⁸
1.5×10⁸
1×10⁸
5×10⁷
0
Tota Red object Integrated Intensity (GCU x µm²/Image)
2.5×10⁵
2×10⁵
1.5×10⁵
1×10⁵
5×10⁴
0
[µM] 0 0.1 1 10 100 500
b
Vehicule
1µM MK8722
TMEM70 WT/WT
TMEM70 KO/KO
TMEM70 SNP/SNP
TMEM70 KO/KO
TMEM70 SNP/SNP
Memb [kDa]
10000 1000 100
Complex I
NDUFV1
NDUFV2
NDUFS1
NDUFA2
NDUFS4
NDUFV3
NDUFA12
NDUFS6
NDUFA7
NDUFAB1
NDUFA6
NDUFA9
NDUFA5
NDUFS2
NDUFS3
NDUFS7
NDUFS8
MT-ND1
NDUFA3
NDUFA8
NDUFA13
NDUFA1
MT-ND6
MT-ND3
MT-ND2
NDUFC2
NDUFA10
NDUFS5
NDUFA11
MT-ND4
NDUFB6
NDUFB5
NDUFB10
NDUFB11
NDUFB1
NDUFB4
MT-ND5
NDUFB7
NDUFB8
NDUFB2
NDUFB3
NDUFB9
NDUFAB1
Complex II
SDHA
SDHB
SDHC
SDHD
Complex III
MT-CYB
CYC1
UQCRFS1
UQCRC1
UQCRC2
UQCRH
UQCRB
UQCRQ
UQCR10
UQCR11
Complex IV
MT-CO1
MT-CO2
MT-CO3
COX4I1
COX5A
COX5B
COX6A1
COX6B1
COX6C
COX7A1
COX7A2
COX7C
NDUFA4
COX7A2L
Complex V
ATP5F1A
ATP5F1B
ATP5F1C
ATP5F1D
ATP5MC1
MT-ATP6
MT-ATP8
ATP5MJ
ATP5MG
ATP5MF
ATP5ME
ATP5MK
ATP5PO
ATP5PD
ATP5PB
ATP5PF
0 0.5 1.0

**Extended Data Figure 6. MK8722 dose-response cytotoxicity and complexome profiling reveal preserved Complex V assembly and increased Complex II abundance following chronic AMPK activation.**

(a) Cytotoxicity assessment of MK8722 treated for seven days across a concentration range (0–500 μM) in TMEM70-deficient cardiomyocytes, using Calcein-AM (live, green) and propidium iodide (dead, red) staining. DMSO served as the vehicle control at a concentration of 0 for MK8722. Representative fluorescence images (top) and quantification of total green and red object integrated intensity (bottom; left and right y-axes, respectively). n=4 from one differentiation. Bars show mean ± SEM.

(b) Complexome profiling heatmaps showing the relative abundance of individual ETC complexes subunits across apparent molecular mass (Memb, kDa), comparing vehicle-treated $TMEM70^{WT/WT}$, $TMEM70^{KO/KO}$, and $TMEM70^{SNP/SNP}$ iPSC-CMs with MK8722-treated $TMEM70^{KO/KO}$ and $TMEM70^{SNP/SNP}$ iPSC-CMs. Color scale indicates relative abundance (0–1).

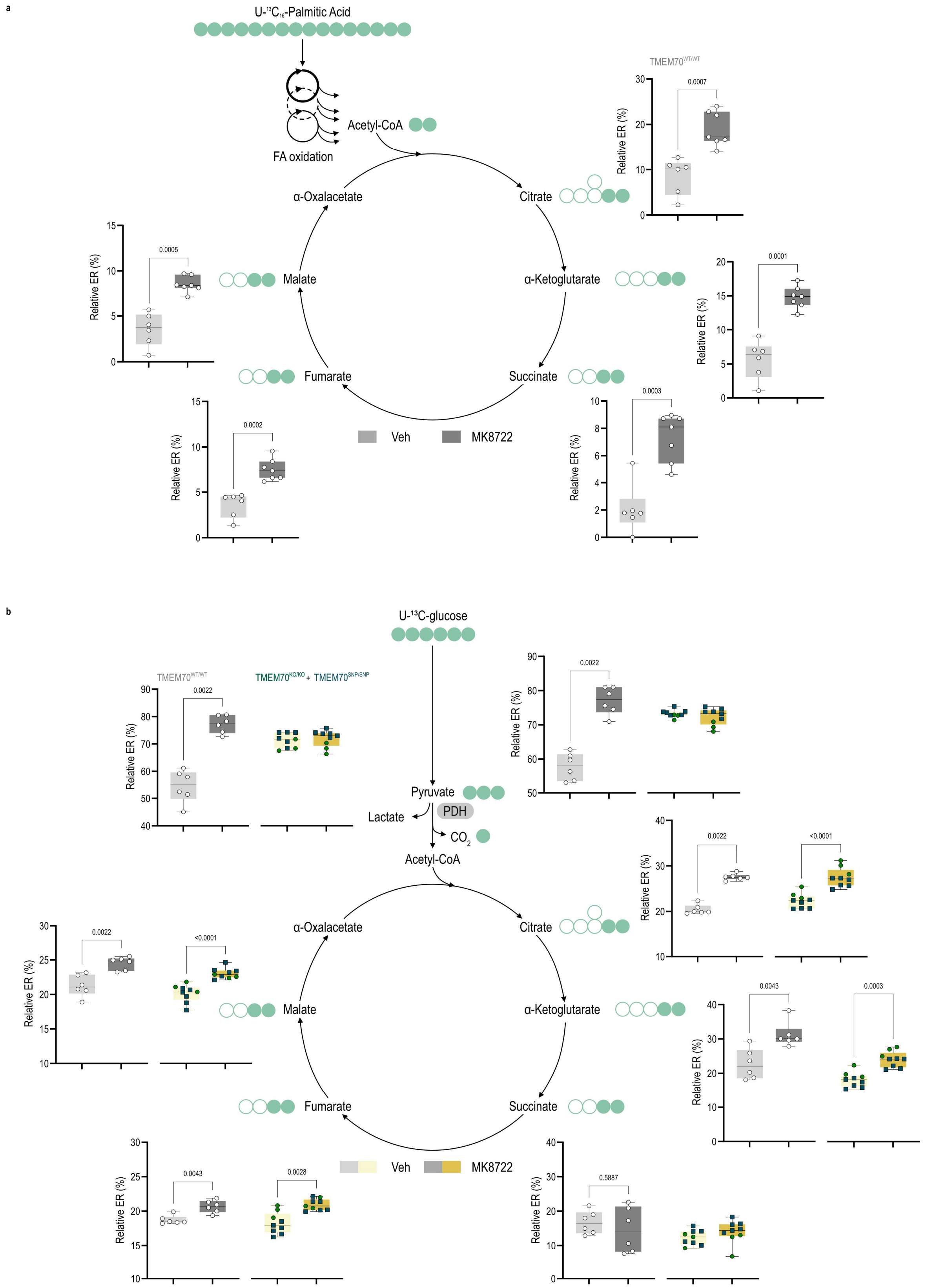


**Extended Data Figure 7. U-$^{13}C_6$ metabolic flux analysis reveals AMPK-driven enhancement of fatty-acid and glucose-derived carbon entry into the TCA cycle.**

(a) U-$^{13}C_{16}$-palmitic acid tracing in vehicle- and MK8722-treated TMEM70$^{WT/WT}$ iPSC-CMs. The schematic illustrates fatty-acid β-oxidation and the entry of labeled acetyl-CoA into the TCA cycle. Relative isotopic enrichment (ER, %) of citrate, α-ketoglutarate, succinate, fumarate, and malate is shown; filled circles indicate $^{13}C$-labeled carbons; n=6 from two independent differentiations. P values were determined by a two-tailed Mann Whitney test.

(b) U-$^{13}C$-glucose tracing in vehicle- and MK8722-treated TMEM70$^{WT/WT}$ and pooled TMEM70-deficient iPSC-CMs. The schematic illustrates glycolytic conversion of glucose to pyruvate, and the entry of labeled acetyl-CoA into the TCA cycle. Relative isotopic enrichment (%) of pyruvate, lactate, citrate, α-ketoglutarate, succinate, fumarate, and malate is shown for each genotype and treatment condition; filled circles indicate $^{13}C$-labeled carbons. TMEM70$^{WT/WT}$; n=12 from two independent differentiations, and *TMEM70*-deficient group; n=9 from three independent differentiations. P values from two-tailed Mann-Whitney test, as indicated.

**Supplemental table 1. Antibody list**

| Primary antibody | Company | Identifier | Assay | Dilution |
|---|---|---|---|---|
| Mouse monoclonal anti TOMM20 antibody [4F3] | Abcam | Cat# ab56783; AB_945896 | Wb | 1 to 1000 |
| Rabbit policolnal anti Phospho-CaMKII alpha/delta (Thr286) | Thermo Fisher Scientific | Cat# PA5-36838; AB_2553769 | Wb | 1 to 1000 |
| Rabbit policlonal anti CaMKII delta | Thermo Fisher Scientific | Cat# PA5-22168, AB_11153337 | Wb | 1 to 1000 |
| Rabbit polyclonal antiTroponin T (Cardiac)TNNT2 Antibody | Cell Signaling Technology | Cat# 5593; AB_10694559 | Wb | 1 to 1000 |
| Rabbit polyclonal anti Ryanodine receptor 2 (RYR2) pSER2814 | Babrilla | Cat# A010-31AP; AB_3665192 | Wb | 1 to 1000 |
| Rabbit polyclonal anti Ryanodine receptor 2 | Sigma | Cat# HPA020028; AB_1856528 | Wb | 1 to 15000 |
| Rabbit polyclonal anti Phospholamban (PLN, PLB) (pTHR17) | Babrilla | Cat# Cat# A010-13AP; AB_3665193 | Wb | 1to 1000 |
| Mouse monoclonal anti Phospholamban (2D12) | Thermo Fisher Scientific | Cat# MA3-922; AB_2252716 | Wb | 1 to 10000 |
| Mouse monoclonal anti SERCA2 ATPase (2A7-A1) | Thermo Fisher Scientific | Cat# MA3-919; AB_325502 | Wb | 1 to 200000 |
| Mouse monoclonal anti Vinculin | Sigma-Aldrich | Cat# V9131; AB_477629 | Wb | 1 to 1000 |
| Mouse polyclonal anti TMEM70 | Proteintech | Cat# 20388-1-AP; AB_10694436 | Wb | 1 to 1000 |
| Mouse monoclonal anti OXPHOS protein complexes | Abcam | Cat# ab110411; AB_2756818 | Wb | 1 to 1000 |
| Mouse monoclonal anti ATP5F1A | Abcam | Cat# ab5432; AB_304883 | Wb | 1 to 1000 |
| | | | IF | 1 to 200 |
| Rabbit monoclonal anti Phospho-Acetyl-CoA Carboxylase (Ser79) (D7D11) | Cell Signaling Technology | Cat# 11818; AB_2687505 | Wb | 1 to 1000 |
| Rabbit monoclonal anti Acetyl-CoA Carboxylase (C83B10) | Cell Signaling Technology | Cat# 3676; AB_2219397 | Wb | 1 to 1000 |
| Phospho-Pyruvate Dehydrogenase alpha1 (Ser293) | Cell Signaling Technology | Cat# 31866; AB_2799014 | Wb | 1 to 1000 |
| Rabbit monoclonal anti Pyruvate Dehydrogenase (C54G1) | Cell Signaling Technology | Cat# 3205; AB_2162926 | Wb | 1 to 1000 |
| Mouse monoclonal anti TRA-1-60 | Abcam | Cat# ab16288; AB_778563 | Wb | 1 to 1000 |
| | | | IF | 1 to 500 |
| Rabbit monoclonal anti Anit OCT4 | Abcam | Cat# ab181557; AB_2687916 | IF | 1 to 250 |
| Mouse monoclonal anti Optic atrophy 1 (OPA1) | Cell Signaling Technology | Cat# 80471; AB_2734117 | Wb | 1 to 1000 |
| Rabbit monoclonal anti Mitofusin-1 MFN1 | Cell Signaling Technology | Cat# 14739; AB_2744531 | Wb | 1 to 1000 |
| Rabbit monoclonal anti Mitofusin-2 MFN2 | Cell Signaling Technology | Cat# 11925; AB_2750893 | Wb | 1 to 1000 |
| Mouse monoclonal anti Mitofilina (MIC60) | Abcam | Cat# ab137057; AB_3676556 | IF | 1 to 100 |

| Secondary antibody | Company | Identifier | Assay | Dilution |
|---|---|---|---|---|
| Horse anti-mouse IgG polyclonal antibody (HRP) | Cell Signaling Technology | Cat# 7076; AB_330924 | Wb | 1 to 5000 |

| | | | | |
|---|---|---|---|---|
| Horse anti-rabbit IgG polyclonal antibody (HRP) | Cell Signaling Technology | Cat# 7074; AB_2099233 | Wb | 1 to 5000 |
| Goat anti-rabbit IgG, Alexa 488 | Thermo Fisher Scientific | Cat# A-11034; AB_2576217 | IF | 1 to 200 |
| Goat anti-mouse IgG, Alexa 546 | Thermo Fisher Scientific | Cat# A-11003; AB_2534071 | IF | 1 to 200 |
| Goat anti-mouse IgG, Alexa 594 | Thermo Fisher Scientific | Cat# A-11005, AB_2534073 | IF | 1 to 100 |

### Supplemental table 2. Nanostring nCounter Elements TagSet panel

| Gene | Accession number |
|---|---|
| GAPDH | NM_001256799.1 |
| POL2RA | NM_000937.2 |
| TBP | NM_001172085.1 |
| HPRT | NM_000194.1 |
| NKX2-5 | NM_004387.3 |
| NR2F2 | NM_021005.2 |
| GATA4 | NM_002052.3 |
| HEY1 | NM_012258.3 |
| HEY2 | NM_012259.2 |
| HCN4 | NM_005477.2 |
| TNNT2 | NM_001276346.1 |
| TNNI1 | NM_003281.3 |
| TNNI3 | NM_000363.4 |
| TTN | NM_133432.1 |
| TTN-N2B | NM_003319.4 |
| TTN-N2BA | NM_001256850.1 |
| MYH6 | NM_002471.3 |
| MYH7 | NM_000257.2 |
| MYL7 | NM_021223.2 |
| MYL2 | NM_000432.3 |
| RYR2 | NM_001035.2 |
| CASQ2 | NM_001232.3 |
| ATP2A2 | NM_001681.3 |
| PLN | NM_002667.3 |
| SLC8A1 | NM_021097.1 |
| SCN5A | NM_198056.2 |
| CACNA1C | NM_199460.2 |
| KCNA5 | NM_002234.2 |
| KCNQ1 | NM_181798.1 |
| KCNH2 | NM_172057.2 |
| NPPA | NM_006172.2 |
| NPPB | NM_002521.2 |
| MEF2C | NM_002397.3 |
| MEF2D | NM_001271629.1 |
| RBM20 | NM_001134363.1 |
| MYBPC3 | NM_000256.3 |

| | |
|---|---|
| NR4A1 | NM_173157.1 |
| ADRB1 | NM_000684.1 |
| ADRB2 | NM_000024.3 |
| CAMK2A | NM_171825.1 |
| CAMK2D | NM_172127.1 |
| ATP5F1A | NM_001001937.1 |
| SLC2A1 | NM_006516.2 |
| SLC2A4 | NM_001042.2 |
| SLC27A1 | NM_198580.1 |
| MYOM1 | NM_003803.3 |
| CRYAB | NM_001885.1 |
| CKM | NM_001824.4 |
| GATA6 | NM_005257.3 |
| NFATC1 | NM_172389.1 |
| NFATC3 | NM_004555.2 |
